# Flexible contextual modulation of naturalistic texture perception in peripheral vision

**DOI:** 10.1101/2020.01.24.918813

**Authors:** Daniel Herrera-Esposito, Ruben Coen-Cagli, Leonel Gomez-Sena

## Abstract

Peripheral vision comprises most of our visual field, and is essential in guiding visual behavior. Its characteristic capabilities and limitations, which distinguish it from foveal vision, have been explained by the most influential theory of peripheral vision as the product of representing the visual input using summary-statistics. Despite its success, this account may provide a limited understanding of peripheral vision, because it neglects processes of perceptual grouping and segmentation. To test this hypothesis, we studied how contextual modulation, namely the modulation of the perception of a stimulus by its surrounds, interacts with segmentation in human peripheral vision. We used naturalistic textures, which are directly related to summary-statistics representations. We show that segmentation cues affect contextual modulation, and that this is not captured by our implementation of the summary-statistics model. We then characterize the effects of different texture statistics on contextual modulation, providing guidance for extending the model, as well as for probing neural mechanisms of peripheral vision.

## Introduction

Central and peripheral vision fulfill different roles in visual perception, as reflected by their different information processing capabilities. The most influential model of peripheral visual processing is the summary statistics (SS) model (Rosenholtz, 2016; Freeman & Simoncelli, 2011; Balas, Nakano, & Rosenholtz, 2009; Parkes, Lund, Angelucci, Solomon, & Morgan, 2001), which proposes that the peripheral visual input is represented using SS of the activations of feature detectors (Figure 1), computed over pre-specified regions of the visual field (termed pooling windows) whose size scales linearly with eccentricity. This model fits in the descriptive paradigm of vision as a hierarchical feedforward cascade of visual feature detectors (Riesenhuber & Poggio, 1999; Doerig et al., 2019), and it is theoretically appealing because replacing a detailed representation of the visual input with a SS results in a significant compression of the visual input. Furthermore, this compression results in a loss of information that could parsimoniously explain the limitations of peripheral vision (Rosenholtz, 2016), including the impairment of target identification by surrounding stimuli (visual crowding (Balas et al., 2009), often regarded as the most important factor in peripheral vision), as well as phenomena related to visual search (Rosenholtz, Huang, Raj, Balas, & Ilie, 2012), scene perception (Ehinger & Rosenholtz, 2016; Freeman & Simoncelli, 2011), and subjective aspects of visual experience (Cohen, Dennett, & Kanwisher, 2016). The SS framework has also been used to explain auditory perception of sound texture (McDermott & Simoncelli, 2011), suggesting a more general role of SS representations.

**Figure 1:**
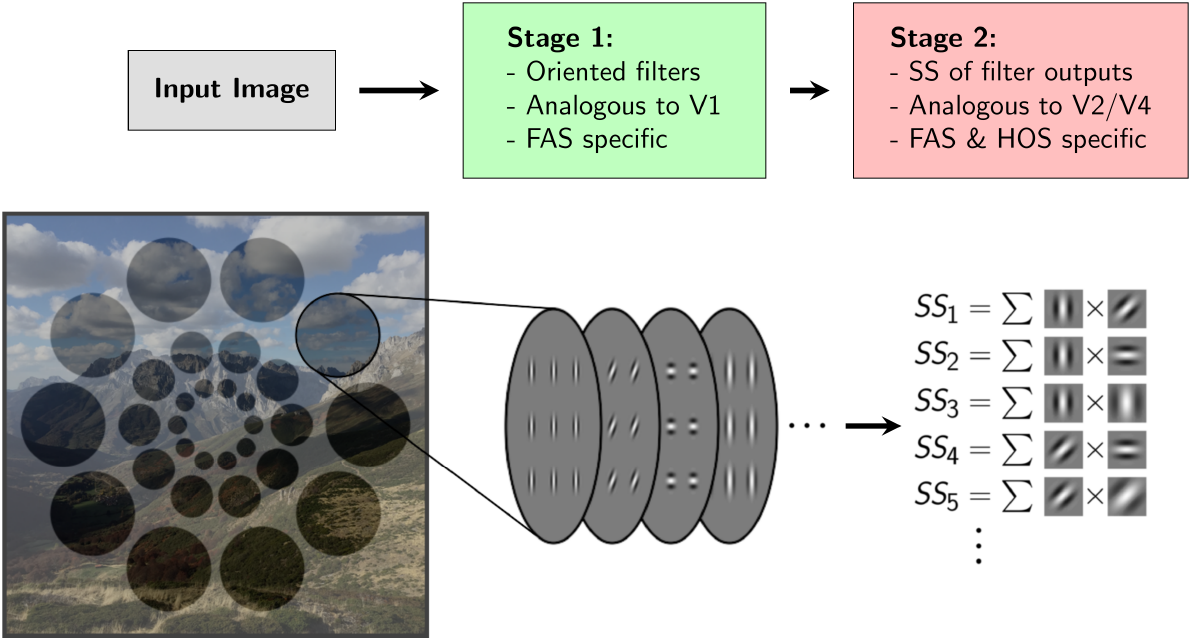
Summary-statistics representation model. Illustration of the main features of the standard SS model, and its relation to physiology and image properties. An input image is first filtered with a bank of oriented V1-like filters, whose activation power is determined by the Fourier amplitude spectrum (FAS) of the image in the pooling region. Then SS are computed over the activations of these filters in fixed pooling windows that tile the visual field. The SS in the second stage are referred to as higher-order statistics (HOS) (in contrast to the statistics contained in the FAS).

Despite providing a solid foundation, it has been hypothesized that phenomena involving segmentation and grouping in peripheral vision escape the standard SS model, and therefore more accurate models of peripheral vision should include recurrent processes of grouping and segmentation (Doerig et al., 2019; Manassi, Sayim, & Herzog, 2013; Manassi, Lonchampt, Clarke, & Herzog, 2016). Grouping different elements into objects or ensembles, or conversely segmenting the scene into different segments, is an essential aspect of human vision. Segmentation processes have been shown to affect several contextual modulation phenomena (i.e. phenomena in which perception of an image region is affected by its surrounds), such as backwards contrast masking (Saarela & Herzog, 2009), the tilt-illusion (Qiu, Kersten, & Olman, 2013), filling-in (Stürzel & Spillmann, 2001; Paradiso & Nakayama, 1991), perceptual fading (Vergeer & van Lier, 2007) and crowding (see (Herzog, Sayim, Chicherov, & Manassi, 2015) for a review). Similar effects have been reported in audition (McWalter & McDermott, 2018; Oberfeld & Stahn, 2012) and touch (Overvliet & Sayim, 2016). In particular, much work with vernier and letter stimuli showed that even small changes to the contextual stimuli, or changes far away from the target, can lead to target-surround ungrouping and a considerable reduction in crowding (Kooi, Toet, Tripathy, & Levi, 1994; Manassi, Sayim, & Herzog, 2012; Manassi et al., 2016; Manassi, Hermens, Francis, & Herzog, 2015; Manassi et al., 2013; Saarela, Sayim, Westheimer, & Herzog, 2009), a phenomenon known as “uncrowding”. It has been argued that these results show a failure of feedforward pooling models, such as the SS model, and that this failure is due to their lack of recurrent processes of grouping and segmentation (Doerig et al., 2019; Doerig, Bornet, Choung, & Herzog, 2020; Herzog et al., 2015; Francis, Manassi, & Herzog, 2017). Furthermore, current SS model implementations also fail to capture the peripheral appearance of natural scenes that contain strong grouping and segmentation cues (Wallis et al., 2019). However, it has been proposed that the SS model may be able to account for these results, without recurrent segmentation or grouping mechanisms that modify the encoding encoding of SS, because segmentation cues could be directly decoded from the fixed SS representation (Rosenholtz, Yu, & Keshvari, 2019). One challenge in exploring these alternatives is that commonly used crowding tasks such as discriminating the inclination of a crowded vernier stimulus (Doerig et al., 2019; Manassi et al., 2013), or discriminating complex feature arrangements in scene distortions (Wallis et al., 2019) often depend on object-specific features, or complex arrangements of features that are not easy to link intuitively or computationally to the more distributed and texture-like representations of the SS model.

Here we test more directly the hypothesis that the SS model does not fully capture segmentation effects on contextual modulation, using naturalistic visual textures which are more easily linked to SS representations. SS representations have long been studied in relation to texture perception, because textures are statistically defined stimuli to texture perception (B. Julesz, 1962; Bela Julesz & Caelli, 1979; Victor, 1994) (the SS model is also referred to as the texture-tiling model of vision (Doerig et al., 2019)). We use naturalistic Portilla-Simoncelli (PS) textures (Portilla & Simoncelli, 2000), which have been instrumental to the recent success of the SS model (Rosenholtz et al., 2012; Ehinger & Rosenholtz, 2016; Freeman & Simoncelli, 2011; Balas et al., 2009) and are a useful experimental tool for probing the model. PS textures are defined by a set of SS that are inspired in natural image statistics and early human vision, and which are the basis of the main implementation of the SS model of peripheral vision. This makes it possible to compare directly perception of PS textures to SS model predictions. Furthermore, it has been shown that, different from primary visual cortex (V1), neurons in higher cortical areas V2-V4 are selective for PS statistics (Freeman, Ziemba, Heeger, Simoncelli, & Movshon, 2013; Ziemba, Freeman, Movshon, & Simoncelli, 2016; Okazawa, Tajima, & Komatsu, 2015; 2017), offering a framework to relate the SS model and peripheral vision to neural mechanisms. Yet, no studies have addressed how peripheral naturalistic texture perception is affected by contextual modulation and by segmentation cues (see (Meinecke and Kehrer, 1994; Schade and Meinecke, 2011; 2009; Morikawa, 2000; Victor, Thengone, and Conte, 2013) for examples with artificial stimuli, and (Wallis and Peter J. Bex, 2012) for a study with natural images that does not explore segmentation).

Therefore, we use a PS texture discrimination task to study contextual modulation and segmentation in peripheral vision within the framework of the SS model. We evaluate how different texture surrounds affect texture perception, and study the influences of grouping and segmentation cues and of surround structure, as well as the relation between this contextual modulation and crowding.

Our results reveal an important role of segmentation processes in peripheral perception of naturalistic texture and highlight limitations of the feedforward framework of visual processing. Furthermore, we link our results to existing versions of the SS model and to previous work on the physiology of the early visual system, pointing to possible computational processes that may underlie the results. Our work can provide guidance for implementing and testing extensions of the standard SS model that include segmentation and grouping.

## Methods

### Participants

A total of 98 adult individual participants (including the authors DH and LG, denoted in the figures by colors blue and green respectively), participated in the experiments, of which 34 were women. All participants had normal or corrected to normal vision.

This study was conducted in accordance with the Declaration of Helsinki and was approved by the Research Ethics Committee of the Faculty of Psychology of the Universidad de la República. Participants gave signed consent to participate in the experiment, and to have the anonymized data from the experiments made available online. Participants were given no economic or course credit reward for their participation in the experiment.

### Texture synthesis

We synthesized grayscale naturalistic textures using the Portilla-Simoncelli (PS) texture synthesis algorithm (Portilla & Simoncelli, 2000) in Octave (John W. Eaton, David Bateman, Soren Hauberg, & Rik Wehbring, 2015). The algorithm first computes a set of statistics over an input image, including mean luminance, contrast and higher-order moments of the pixel histogram; and the means and pairwise correlations of the activations of multi-scale, multi-orientation filters (steerable pyramid (Simoncelli, Freeman, Adelson, & Heeger, 1992)) analogous to V1 cells. Then it iteratively modifies a white noise image until its statistics match those of the input image. We used as input images natural textures from the Brodatz texture database, the Amsterdam Library of Textures (Burghouts & Geusebroek, 2009) and from the database presented in (Lazebnik, Schmid, and Ponce, 2005). We refer to an image synthesized this way as a naturalistic texture or PS texture. We used filters with 4 scales and 4 orientations, and 9 by 9 pixels neighborhood (corresponding to a 0.3° x 0.3° neighborhood with the viewing distance used) for computing the spatial correlations of the filter responses. We synthesized two 1024 x 1024 PS textures for each input image.

For each PS texture we also synthesized a phase-scrambled texture. This was achieved by first generating a uniform noise image and then replacing its Fourier amplitude spectrum (FAS) for the FAS of the naturalistic texture. Thus, this procedure produces a pair of PS textures and a pair of phase-scrambled textures that are used in the experiments.

Phase-scrambling a naturalistic image can change the histogram of pixel activations (e.g. changing the minimum and maximum intensities). To prevent participants from using aspects of the pixel histogram (e.g. brightness) as cues to solve the task, we matched the pixel histograms of the naturalistic and phase-scrambled images to an average of the two, using the SHINE package for Octave (Willenbockel et al., 2010) with 30 iterations. In each iteration their FAS was also matched to the original FAS, and the structural similarity index (SSIM) with respect to the original image was also optimized in order to reduce alterations to image structure (Zhou Wang, A.C. Bovik, H.R. Sheikh, & E.P. Simoncelli, 2004; Willenbockel et al., 2010). Images produced by this method appeared very similar to the starting textures (besides changes in pixel intensities), suggesting it did not produce noticeable structural alterations.

In experiment 3, to generate the surround image that was dissimilar to the target only in HOS, we started by generating a new PS texture using a different input image than the one used for the target. Then we matched its FAS and pixel histogram to those of the target PS texture with the SHINE package, using 30 iterations. In each iteration the SSIM with respect to the original surround PS texture was also optimized. For the surround texture that was dissimilar in both FAS and HOS, the same procedure was used but without matching the FAS to the target PS texture.

### Texture selection

Since there is considerable variation in the discriminability of different PS textures from their phase-scrambled counterparts (Freeman et al., 2013), we synthesized a large set of pairs of PS and phase-scrambled textures and selected those that subjectively appeared to have high discriminability, to make the task easier. We also selected textures that had different kinds of structures, in order to better probe the texture space (e.g. strongly oriented, weakly oriented, regular, irregular).

Also, in experiment 3 most textures to which we applied the FAS matching procedure acquired a phase-scrambled appearance, so we selected for further use those that maintained a naturalistic appearance after this procedure.

Due to resource contstraints and design choices, we did not use the same number of textures for each experiment. The textures used in each experiment are those shown in the corresponding figure.

### Organization of experimental sessions

An experimental session consists of one participant performing an experiment with a given texture. When a participant performed an experiment with more than one texture, these were used separately in different experimental sessions. No participant performed the same experiment twice with the same texture. Not all participants in a given experiment performed the same number of experimental sessions (i.e. some participants performed the experiment with more textures than others). For each experiment we report the total number of experimental sessions, corresponding to the sum of the experimental sessions performed by all participants. In total, participants completed 189 experimental sessions across all experiments

Experiment sessions were divided into 2 to 4 experimental blocks (with balanced conditions) separated by 30 s resting periods. Within these blocks, some conditions were also blocked (with their order balanced across participants) to avoid confusions during the task. The different conditions contained within a block were randomly interleaved and each was presented an equal amount of times. The total duration of the experiments, including training and instructions, was between 20 and 45 minutes.

Sometimes participants performed two experimental sessions one after the other, using two different textures for a given experiment.

### Stimulus sampling

All textures shown in the experiments were patches cropped from these larger synthesized images, with a linear transparency gradient at their border, allowing for a smooth fading with their neighboring surface e.g. the background or a neighboring texture). These gradients had a length of 4 pixels, roughly equivalent to 0.15°. For each texture patch displayed, the cropped region was randomly selected over the whole image on a trial by trial basis.

We note that because the PS statistics were matched over the large synthesized images, the random sampling of patches from these images introduced some trial by trial variation in the texture statistics displayed. Although testing the effect of this image variability on our results would require additional experiments, we think this variability is unlikely to have significant effects on the participants performance as discussed in section S6.

In each individual trial an angle multiple of 90° was randomly chosen and all textures were rotated by this angle before being cropped for display. This was done to reduce participants adaptation to low level properties of the textures.

### Task

Our task is a variation of that described by Freeman et.al. (2013), and consisted in discriminating between the naturalistic and the phase-scrambled versions of a texture.

The target stimuli (targets) consisted of two circular patches of texture presented simultaneously for 233ms, centered at 12° to the right and to the left of the fixation point (Figure 2). We used three different target configurations: 1) phase-scrambled target to the right (PS texture to the left), 2) phase-scrambled target to the left (PS texture to the right), or 3) no phase-scrambled target (PS texture in both targets). The three configurations were shown an equal amount of times, in random order. Participants were instructed to report the location of the phase-scrambled target with the arrow keys, and to use the upwards arrow to indicate the absence of phase-scrambled targets. This task design with two targets and 3 conditions was used to discourage participants from looking away from fixation, to compensate for the lack of eye-tracking in the experiments.

**Figure 2:**
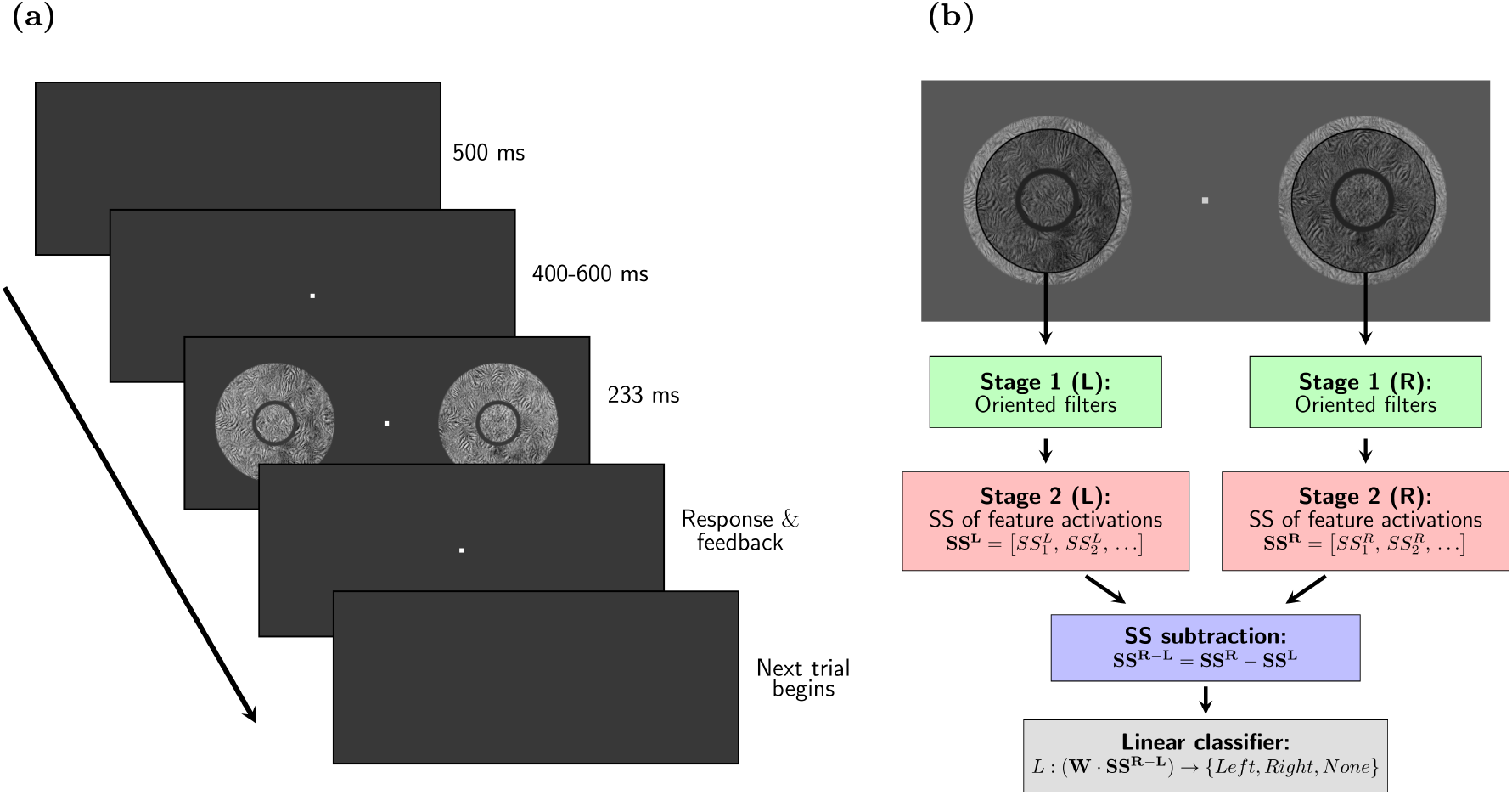
Task design and observer model. **(a)** Two targets centered at 12° to the left and to the right of the fixation point were displayed simultaneously in each trial for 233 ms. Either the left, the right, or none of the targets was sampled from the phase-scrambled texture (with the others sampled from the naturalistic texture), and the participant had to indicate with the arrows where (if) the scrambled texture was present (3 AFC). In most trials, we added uninformative surround textures around the target (in this example surround textures are present, separated from the target by a gap). In any given trial, the two targets always had the same kind of surround. To aid visibility, the size and color of the fixation dot in this image are not the same as in the experiments. **(b)** Diagram showing the architecture of the model observers based on the SS model used to simulate the experiments. The SS of the PS model are computed over circular pooling windows centered on each target (illustrated by the shaded regions). The difference between the SS of the two targets is used to predict the stimulus configuration (i.e. where the phase scrambled target texture is). See Methods for implementation details.

The sequence of events in any given trial was the following (see Figure 2): 1) Start with the gray screen, 2) after 500 ms a red fixation dot appeared at the center of the screen, which participants were instructed to fixate, 3) after a time interval sampled uniformly from 400 ms to 600 ms, the two targets were presented simultaneously for 233 ms (14 frames), 4) after the targets disappeared the participant responded (without a time limit), 5) auditory feedback was provided and the fixation dot disappeared, returning to step 1). Participants were told to use the response stage (step 4) to rest as needed by delaying the response.

For experiment 4 we slightly modified the task for half of the participants. In this variation of the task, participants were instructed to indicate the position of the PS texture, instead of the phase-scrambled texture. Accordingly, we substituted the condition with two naturalistic targets for a condition with two phase-scrambled targets, this maintaining the structure of the task.

### Surround textures

In all experiments we included surrounding textures with varying shapes and texture contents. In any given trial the two targets shared the same kind of surround. These surrounds were also sampled randomly (and independently from each other and from the targets) from the larger synthesized textures. Unless indicated otherwise in the text, the surrounds were sampled from the PS texture of the texture pair to be discriminated in the targets.

In most cases, surrounds were rings (or half-rings) with a width (i.e. distance between inner and outer edges) equal to target diameter. Experiment 1 and texture T1 in experiment 3 were an exception, having a surround width 1.4 times the diameter of the target. The surrounds of the split-disk targets in experiment 2 were not rings, but they had the same outer diameter as the surrounds for the corresponding disk-shaped targets.

Surrounds could be contiguous to the target or separated by a gap showing the gray background. The gap had a width of 0.5° in all cases except experiment 1, where it had a width of 0.35° and texture T1 in experiment 3, where both gaps of 0.5° and 1° were used (although these were grouped together for the analysis, see Supplementary S5). We selected this gap width by subjective visual inspection, considering the need for a gap large enough to be clearly visible in the periphery, but as small as possible to minimize the spatial differences between the stimuli with and without a gap (see Supplementary S2 and S4 for more information on the slight variability in gap size in some conditions).

### Training and difficulty adjustment

Before the experiment, participants were provided with training opportunity. Auditory feedback was used in all stages of training, as well as in the main experiment. In the first training session, targets were shown without surround and remained on the screen until the participant responded. The second training session also used targets only, but had the same dynamics as the experiment. Both sessions were terminated at will by the participant by pressing a special key.

After running experiments 1 and 3 with texture T1 with a target diameter of 3.5°, we observed considerable variability between participants in task performance. Therefore we adjusted task difficulty to each participant (except when noted otherwise), in order to drive participants to a more informative performance range (preventing saturation with very high or with chance-level performances). To this aim, we presented a sequence of trials with unsurrounded targets in which target diameter was adaptively adjusted using the accelerated stochastic approximation procedure (Treutwein, 1995) to drive participant performance to a predetermined level of 90% correct responses (see Supplementary S7 for details on the procedure and final sizes distribution). If the final target diameter was larger than 5.3° (160 pixels), we used a diameter of 5.3° in the experiments. The widths of the surrounds were then set equal to the target diameter. We note that the results from experiment 1, and experiment 3 with texture T1 were obtained without size adjustment, because this procedure was only incorporated after these experiments.

After size adjustment, we repeated the static and dynamic training stages as described above, including also the surrounds, and instructed participants to perform the task ignoring the surrounds. Again, participants terminated these sessions at will.

### Materials and apparatus

The task was performed in a dark room, using a 27 inch LCD screen (ASUS, model PG278QR) with a refresh rate of 60 Hz. Participants used a chinrest to maintain a viewing distance of 40 cm, at which 1° of the visual field subtended 30 pixels. Experiments were ran on Psychtoolbox-3 (Kleiner et al., 2007) running in Octave Version 4.0.0 in Ubuntu 14.04.

The background gray had a luminance of 8.7 cd m^−2^, and the textures used in the experiments had a range of mean luminances of 50.9 cd m^−2^ to 67.3 cd m^−2^, and a range of standard deviations in the luminance of the pixels of 26.9cdm^−2^ to 34.1 cdm^−2^, as determined with a screen calibration performed with a colorimeter (Cambridge Research Systems, model ColorCAL II).

### Summary-statistics model observer

We implemented an image-computable observer model based on the feedforward SS model with fixed pooling windows (Freeman et al., 2013). This model first computes PS statistics over the two stimuli, then computes their difference and feeds it to a linear classifier to solve the task (see Figure 2b). The weights of the discriminator were optimized to maximize discrimination performance on a training set, and the model is then tested on a separate test set (cross-validation). We added noise to the PS statistics computed by the model in both training and testing stages, to roughly match the performance of the human participants on average across stimuli.

We first generated sample images of single stimuli such as those used in the experiments, with either phase-scrambled or naturalistic targets diameter 110 pixels (corresponding to a diameter of 3.7° in the experiments), and with the different surrounds. We adapted the code of (Freeman & Simoncelli, 2011) to compute PS statistics over a circular fixed pooling area centered on the target. We used a pooling area with a diameter of 360 pixels, equivalent to 12° of visual field. We based this pooling size on Bouma’s law of crowding (Whitney & Levi, 2011), which says that surround elements hinder target perception when they are within a distance of about 0.5 times the eccentricity, thus we used this distance (12° × 0.5) as the radius of integration around the target center. We note that previous studies on the SS model (Rosenholtz et al., 2019; Freeman & Simoncelli, 2011; Wallis et al., 2019; Doerig et al., 2019) use multiple pooling regions with smaller sizes (with their diameter and not their radius equal to half the eccentricity, analogous to V2 receptive fields) that tile the visual field. Although such models are more realistic than our model, and their structure may allow them to capture some more complex phenomena, using multiple pooling regions would require a more complex decoder and several additional design choices. Therefore, in the interest of simplicity, we opted for the single pooling window matching Bouma’s law.

We computed PS statistics using 4 scales, 4 orientations, and a neighborhood for computing spatial correlations of 7 pixels (smaller than for texture synthesis to reduce the number of model parameters), corresponding to 0.7° of visual field in the experiments. This procedure leads to 782 SS per stimulus (after removing the repetitions of symmetric parameters from the correlation matrices).

To mimic the experimental task we arranged the stimuli (which either had naturalistic or phase-scrambled target) into 3 kinds of ordered pairs, equivalent to those shown in the experiment. Using *Nat* and *Scr* to refer to stimuli with naturalistic and scrambled targets respectively, the 3 kinds of ordered pairs were: {*Scr, Nat*}, {*Nat, Scr*} or {*Nat, Nat*}. As in the experiment, the stimuli from a given pair had the same surround. Then, we subtracted the SS of the second stimulus to each corresponding SS of the first stimulus, resulting in 782 differences in SS (or predictors) for each stimulus pair. The observer consisted of a linear discriminator trained to predict the class of the stimulus pair (e.g. {*Scr, Nat*}, {*Nat, Scr*} or {*Nat, Nat*}) from the SS difference of the pair.

First, for an observer trained for a given experiment, we generated 750 stimulus pairs (250 of each class), or trials, for each different surround condition in the experiment, and computed the difference in SS (predictors) for each generated pair. We then added Gaussian noise to the predictors, with a standard deviation equal to the standard deviation of the predictor across the training dataset containing all the conditions for the simulated experiment). Next, we normalized each predictor to have unit variance (using the default setting of the fitting package, glmnet (Friedman et al., 2019)). Lastly, we trained multiclass logistic regression on the normalized predictors (i.e. the differences in SS with added noise) with L2 penalization, and optimized the hyperparameter that weights the penalization by 10-fold cross-validation (i.e. the default in the glmnet package). For each experiment, we trained 8 different models (observers), using different noise samples and different samples for the training set, leading to some variability between model observers.

After training the models, we tested their discrimination performance on a test set comprising 1500 texture pairs (500 of each class) for each surround condition.

We verified that all the trends and conclusions are robust to the choices of target size, penalization (we tested also elasticnet, which uses a mixed L1 and L2 penalization), and noise level. Furthermore, we also ran the model with a variation of the task that involved no stimulus sampling variability (see Supplementary S6).

### Statistical analysis

All experiments were first performed with texture T1, and all but experiment 1 were then reproduced with other textures. Experiments performed with T1 sometimes had more conditions than experiments with the other textures. These conditions exclusive to T1 are analyzed separately in the supplementary analysis.

We analyzed the data of the experiments and the simulations using generalized linear mixed models (GLMM) of the binomial family (Gelman & Hill, 2006). In these models we included a fixed effect for each parameter of interest and an offset term. For each of the fixed effects we added random effects. When applying the GLMM to multiple textures to estimate the mean effect across textures (e.g. Figure 5), we included for each fixed effect a random effect for texture and a random effect for participants nested within texture. We also applied the GLMM to individual textures, both for the analysis of data that was only collected with one texture (e.g. experiment 1), and for estimating the effects of the different manipulations on each texture. In the plots showing the effects for multiple textures (e.g. Fig. 5), the estimate for each individual texture was obtained by fitting a GLMM to that texture individually. In these cases we only used a random effect for participants. Correlations between random effects in the model were always set to 0, to avoid overly complex models (Bates, Kliegl, Vasishth, & Baayen, 2015).

All the GLMM fitted by maximum likelihood using the R package lme4 (Bates et al., 2019). The reported p-value for each effect was obtained by a likelihood ratio test (LRT) between the full model and the null model, in which that fixed effect is set to 0. The 95% confidence intervals of the fixed effects were obtained by likelihood profiling.

The analysis in the text is based on the parameters fitted by these models, which are in log-odds ratio (LOR) units. Although less intuitive than simple differences between success probabilities, this is a more adequate measure for the experimental effects, especially given the variability in performance between participants and textures.

In some cases we fitted a GLM to the data of each participant in an experiment in order to display the actual observed LOR for each individual (e.g. Figure 3). These models contained no random effects. The confidence intervals for the parameters were obtained by the Wald method, and their p-values by the Wald test.

**Figure 3:**
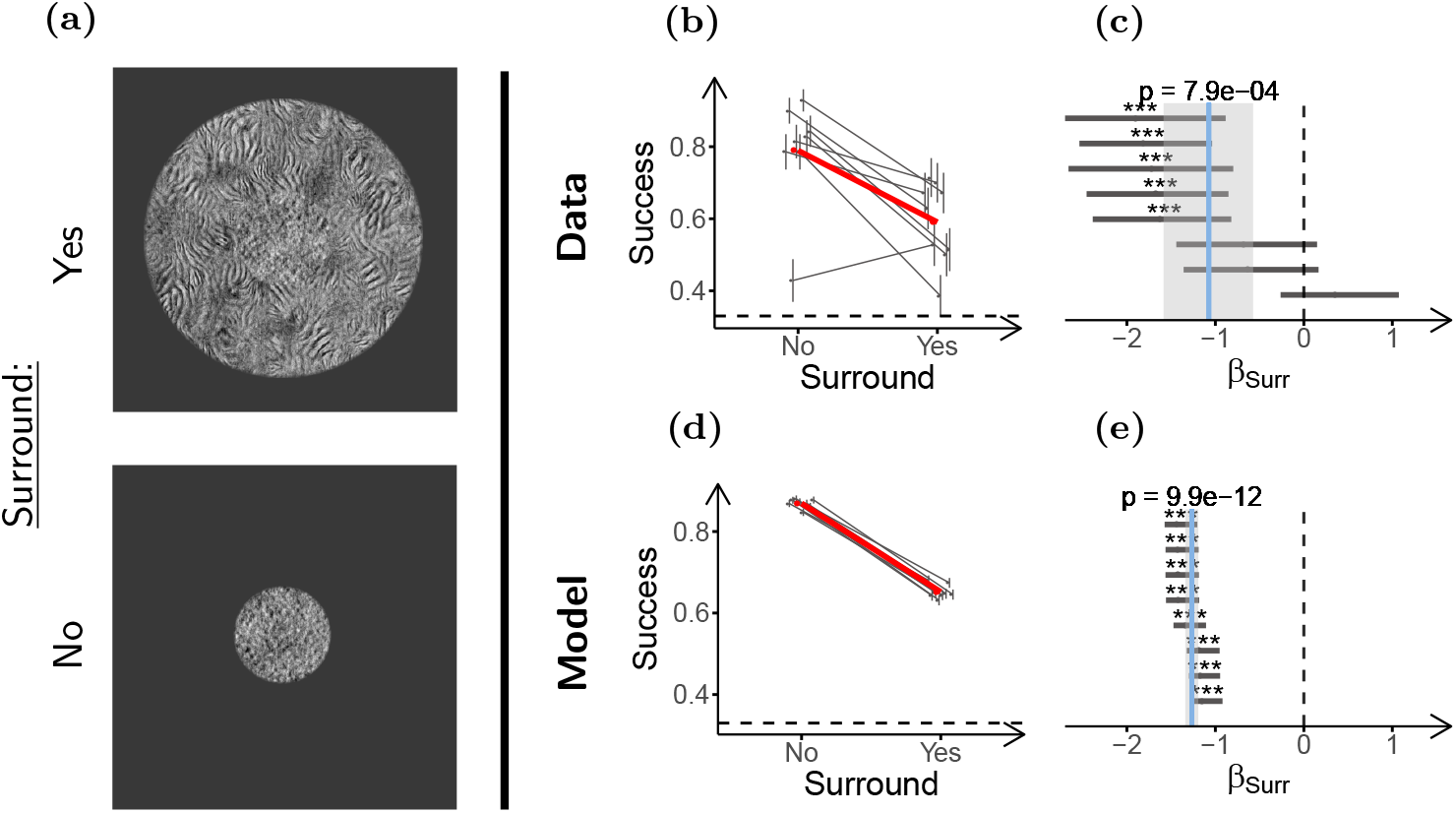
Surround textures impair texture discrimination performance. **(a)** Stimulus configurations used in the experiment (only scrambled targets shown). Top: Target with surround, Bottom: Target without surround. **(b)** Task performances for the two conditions. The gray dots and lines show the performance of individual participants. Vertical lines indicate the ±SD of the estimated performance. Horizontal jitter was applied to aid visualization. The larger red dots show mean performance across participants for each condition. The dashed horizontal line shows chance performance. **(c)** Log-odds-ratios (LOR) between the presence and absence of the surround (*β_surr_*), estimated from the performance data in **(b)**. Each dot shows the LOR for one participant (estimated by fitting a GLM), and the horizontal lines indicate their 95% c.i. Statistical significance of the LOR for the individual participants obtained by the Wald test is indicated as follows: *p* < 0.05 (*), *p* < 0.01 (**), *p* < 0.001 (* * *). The vertical solid blue line indicates the estimated mean LOR for the population (estimated by fitting a GLMM), and the grey shade indicates its 95% c.i. The p-value of the mean LOR estimate as obtained by likelihood-ratio test (LRT) is indicated above the solid line. The dashed vertical line marks the LOR at which there is no difference between the conditions. & show the same as **(b)** & **(c)** but for the model observers. Participants (n=**8**) performed 70 trials in each condition, and model observers (n=8) discriminated 1500 stimuli per condition.

We excluded from analysis experimental sessions in which the participant performed below 45% correct for all conditions (chance level performance is 33%), to avoid strong floor effects. This criterion discarded 14 of the total 189 experimental sessions. In the main text we report for each experiment the number of experimental sessions that satisfied the inclusion criterion. All results and analysis are robust to removing this exclusion criterion, as well as to excluding the main author from the analysis.

Data analysis was performed in R 3.4.4 (R Core Team, 2018) using the packages lme4 1.1-19 (Bates et al., 2019), dplyr 0.7.6 (Wickham, Romain François, Lionel Henry, & Kirill Müller, 2018), tidyr 0.8.1 (Wickham, Lionel Henry, & RStudio, 2018), ggplot2 3.0.0 (Wickham, 2016), broom 0.5.0 (Robinson & Alex Hayes, 2018), MASS 7.3-50 (Venables & Ripley, 2002), and this document was generated using knitr 1.20 (Xie, 2015).

### Data availability

The anonymized raw data of the experiments, together with the analysis code, and the code for running the experiments, are available in the Open Science Framework (https://osf.io/8zr5h/). All participants gave informed written consent for their anonymized data to be publicly shared.

## Results

We used a PS texture discrimination task (details in Figure 2 and Methods) to study contextual modulation of texture perception in peripheral vision. We refer to contextual modulation as the observed phenomenon by which perception of a part of a visual stimulus is affected by its surrounds, regardless of the precise underlying mechanisms. In our experiments we measure changes in contextual modulation as the changes in task performance between conditions with different surrounds (taken as indicative of changes in target perception between conditions induced by these surrounds). PS textures are characterized by a set of SS inspired in natural image statistics and early human vision, including the correlations between the outputs of V1-like filters selective for orientation and spatial frequency. The corresponding SS model implementation consists of two stages: the first stage computes the responses of the V1-like filters to the input image, and the second stage evaluates the PS statistics of those filter activations within fixed pooling windows (Figure 1).

The task required discriminating patches of naturalistic PS texture from their corresponding phase-scrambled textures (see Figure 2) in a 3 alternative forced choice design (we refer to the patches to be discriminated as targets). These PS and phase-scrambled texture pairs have the same Fourier amplitude spectrum (FAS), which means they activate the V1-like filters of the SS model with the same average energy, and are thus matched in the first stage of the SS model. Unlike phase-scrambled textures, PS textures also have a more structured distribution of filters activations, corresponding to higher order statistics (HOS) that drive the second stage of the SS model and lead to a more natural appearance (Portilla & Simoncelli, 2000).

To evaluate whether our experimental observations could be captured by the feedforward SS model with fixed pooling windows, we implemented a model SS observer to solve the task using a linear classifier on the PS statistics of the stimuli, computed over a fixed area centered on the target (Figure 2b, Methods). We then compared qualitatively the model’s discrimination performance to the participants. The radius of the pooling windows was chosen according to Bouma’s law of crowding, which says that surrounding stimuli can interfere with target identification when they are within a distance of approximately 0.5 times the target eccentricity (Pelli, Palomares, & Majaj, 2004) (see Methods).

The results are divided into three sections. First, we report the effect of the surround on performance, and its dependence on target-surround grouping or segmentation. Then we explore the relevance of the statistical structure of the surround texture to contextual modulation. Lastly, we study the relation of this contextual interaction to crowding.

### Contextual modulation and grouping

Target-surround grouping, or conversely segmentation, is a major modulator of contextual interactions in vision, especially for crowding (Manassi et al., 2013; Saarela & Herzog, 2009; Qiu et al., 2013; Levi, 2008). It has been argued that these segmentation and grouping processes are an important missing component in pooling models of peripheral vision, including the SS model (Manassi et al., 2013; Wallis et al., 2019; Doerig et al., 2019). Despite considerable work using stimuli such as objects, shapes or features (e.g. (Manassi et al., 2013; Manassi et al., 2016; Saarela, Westheimer, and Herzog, 2010; Kooi et al., 1994)), our understanding of how grouping processes affect peripheral perception is still incomplete because it is not clear how to relate those tasks that use non-texture stimuli to the SS model, which may be affected by more global stimulus information (Rosenholtz et al., 2019), and whether those results extend to texture processing.

Thus, to better understand the role of grouping and segmentation in the SS model, and how they influence perception of textures, we sought to determine whether contextual modulation of naturalistic texture perception is affected by segmentation or grouping cues.

#### Experiment 1: Target-surround discontinuity reduces contextual modulation

First, we measured whether naturalistic texture perception is affected by contextual modulation. Based on the relevance of contextual modulation for target identification in peripheral vision, we expected task performance to be impaired by surrounding textures. To test this, we presented participants (n=8) with targets in isolation, and with targets surrounded by an uninformative texture ring that was sampled from the same PS texture (Figure 3a).

As expected, task performance was considerably worse for the surrounded targets (Figure 3b). To quantify the effect sizes and test for their statistical significance, we fitted a generalized linear mixed model (GLMM) to the data (which allows to take into account between-participant variability; see Methods and section S1). We report the log-odds ratio between the conditions (LOR, denoted by *β*), which is a measure of their difference in success probability (see a guide for converting between the two in section S1). For example, *β_surr_* quantifies the effect of the surround around the target, and *β_surr_* <0 means that the surround hindered performance. Figure 3c shows that in our experiments the surround strongly impaired performance, and that the effect was statistically significant (*β_Surr_* = −1.07, ci = [−1.58, −0.57], p = 8 × 10^−4^). This effect was captured by our implementation of the SS model (Figure 3e).

We next tested whether segmentation affects this contextual modulation, and whether the effect can be captured by our SS model implementation. To probe the effect of segmentation, we presented participants (n=9) with two kinds of stimuli, either with continuous target and surround, or with a visible gap that induced target surround segmentation (Figure 4a). Importantly, the gap was generated by shrinking the target of the continuous stimuli, keeping surround geometry the same in the two conditions. With this design, if pooling regions are constant, the two conditions would have the same amount of surround texture pooled with the target, but in the discontinuous condition there would be less target texture to be integrated (due to the smaller target size). In line with what could be expected from the the ratio of informative target texture and uninformative surround texture for each stimulus, our implementation of the SS model showed worse performance in the discontinuous than in the continuous condition (Figures 4d, 4e; a similar reasoning to that applied in (Manassi et al., 2013)). This is in contrast to what we expect from previous studies using simple stimuli, in which segmentation reduced contextual modulation (Qiu et al., 2013; Saarela et al., 2010; Kooi et al., 1994; Manassi et al., 2012; 2013; Manassi et al., 2015; Manassi et al., 2016). Figure 4b shows that performance increased moderately when target and surround were discontinuous (*β_Discont_* = 0.62, ci = [0.37, 0.87], p = 1 × 10^−4^, Figure 4c). Thus, our SS model implementation was unable to capture the effect of segmentation (see section S6 for further discussion).

**Figure 4:**
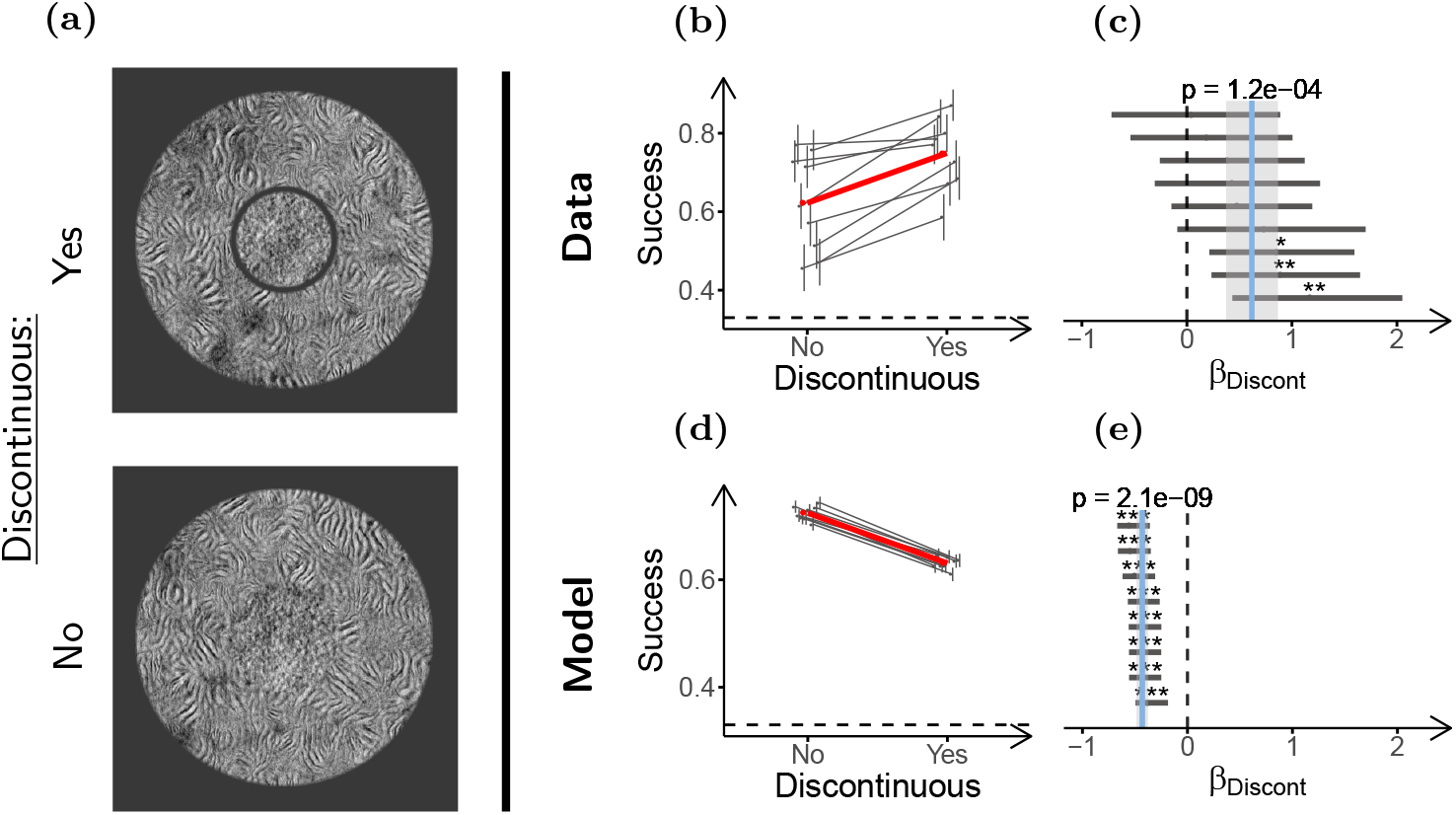
Segmentation reduces contextual modulation. **(a)** Stimulus configurations used in the experiment (only scrambled targets shown). Top: Discontinuous stimulus (smaller target size) Bottom: Continuous stimulus (larger target size). **(b)** Task performance for the two conditions. **(c)** LOR for discontinuity (*β_Discont_*), estimated from the performance data in. & Same as **(b)** & but for the simulated observers. Participants (n=9) performed 70 trials in each condition, and model observers (n=8) discriminated 1500 stimuli per condition. Panels **(b)**-**(e)** use the same conventions as Figure 3.

We also found that the observed effect of discontinuity is sensitive to the size of the gap (see section S2, Figure S2), likely because the gap size affects gap visibility, and also the difference in target sizes between the conditions. Also, notice that segmentation did not completely remove contextual modulation, that is, performance was still lower for the discontinuous surround than for the target alone (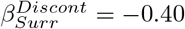, ci = [−0.68, −0.15], p = 5 × 10^−3^, Figure S3).

#### Experiment 2: The effect of target-surround discontinuity is mediated by segmentation

We reasoned that the gap between target and surround used to induce segmentation may also affect performance by other mechanisms, such as reducing the uncertainty of target location within the stimulus, or altering the SS of the stimulus in a way that is not captured by our SS model implementation. For example, it has been proposed that some uncrowding results may be explained by a better encoding or decoding of target information from the SS of the stimuli, allowed by the specific stimulus configurations that generate uncrowding (Rosenholtz et al., 2019). Given that the gap in our stimuli is co-localized with the target, it is possible that their low-level features induce changes in the SS that allow for a better decoding of target information (see Supplementary S6 for modeling results suggesting that such factors may be relevant).

To control for the possible cues related to the gap but not to segmentation, we introduced a different target shape (split-target) consisting of two adjacent semicircles with their straight sides facing outwards (Figure 5a). This split-target shape had approximately the same texture area as the original disk target, and a gap could be introduced around its curved sides, while preserving target-surround continuity on the straight sides of the target.

**Figure 5:**
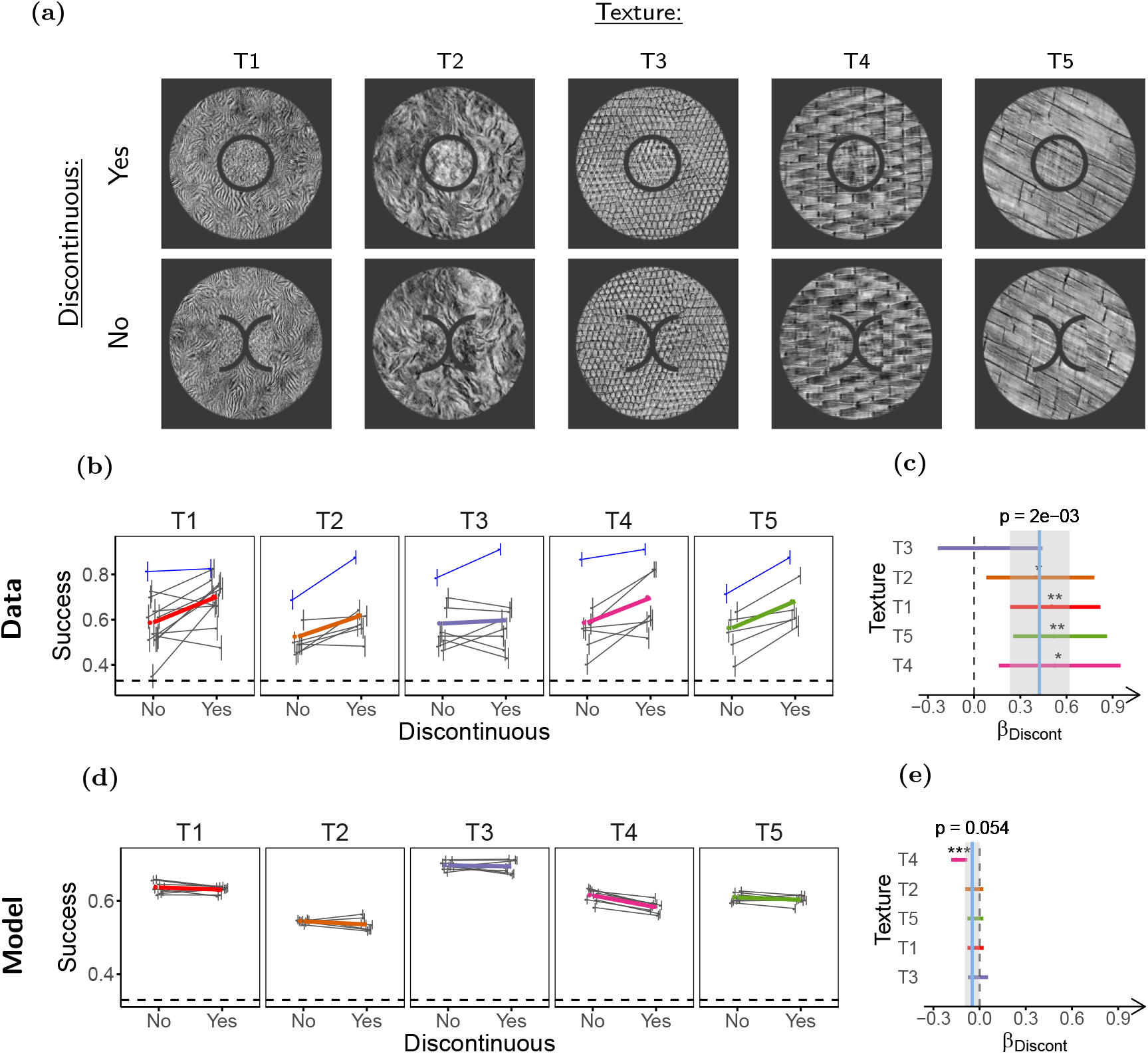
Low level properties of the gap do not explain the effect of discontinuity. **(a)** Stimuli used in the experiment. Top: Disk targets (discontinuous), Bottom: Split-targets (continuous). **(b)** Task performance. Each panel shows the results for a different texture, with texture identity indicated above the panel. The layout of each panel is the same as in 3b, except a different color is used to identify the mean performance for each texture. The data of author DH are indicated by the blue symbols. **(c)** LOR for target-surround discontinuity (*β_Discont_*). The colored dots show the LOR obtained by fitting a GLMM for each individual texture (color coded as in **(b)**) and the horizontal lines indicate their 95% c.i. The p-value for the (*β_Discont_*) of each individual texture, estimated by LRT, is indicated as follows: *p* < 0.05 (*), *p* < 0.01 (**), *p* < 0.001 (***). The vertical solid blue line shows the mean *β_Discont_* across textures and participants estimated by a hierarchical GLMM model using all textures, and the shaded gray region shows its 95% c.i. The p-value for this estimate obtained using LRT is indicated above the line. The dashed vertical line marks the value at which there is no difference between conditions. **(d)** & **(e)** Same as **(b)** & **(c)** but for the model observers. Participants (n=25) completed 40 experimental sessions (see Methods), and performed between 80 and 112 trials per condition. Model observers (n=8) discriminated 1500 trials per condition.

Although the circular targets and the split targets had gaps with similar low level properties, we expected no segmentation for the split-target stimulus because target-surround continuity is maintained. Thus, if the effect observed in the previous section was mediated by segmentation, we should find lower performance for the grouped continuous stimulus (split-target) as compared to the segmented discontinuous stimulus (disk-target). If the effects were mostly due to other factors introduced by the low level properties of the gap, then we would expect similar performance for these two kinds of stimuli.

We presented participants (n=25) with the disk-target and split-target stimuli using 5 different textures to verify that the results did not depend on a specific texture (most participants were shown only some of the textures, see Methods). Participants completed 40 experimental sessions (an experimental session consists of a participant completing the experiment with one texture) that satisfied the inclusion criterion (see Methods). Consistent with a role of segmentation in contextual modulation of texture perception, performance was moderately worse for the continuous (split-target) than for the discontinuous stimulus (*β_Discont_* = 0.42, ci = [0.23, 0.62], p = 2 × 10^−3^, Figure 5c). In contrast to this observation, our implementation of the SS model showed little difference between the stimuli, showing again a failure to capture the segmentation effect (Figure 5e).

We also verified, using additional stimuli for texture T1 (see Figures S4, S5a), that splitting the target had a small and non-significant effect on performance (*β_Split_* = −0.09 ci = [−0.31,0.14], p = 0.43, Figure S5c) validating the use of this experiment to control for low-level gap properties. Furthermore, the estimated effect of the gap after accounting for segmentation was also close to 0 (*β_Gap_* = 0.04, ci = [−0.15,0.22], p = 0.71, Figure S5d), suggesting that effects of the gap other than inducing target-surround segmentation are negligible in our task. This extended analysis supports the interpretation that the effect of segmentation on contextual modulation cannot be wholly explained by the changes in target encoding or decoding allowed by its co-localization with the gap, although it remains possible that a more complex SS model with more statistics or more complex structure could capture these results.

We conclude from these experiments that target-surround segmentation is an important factor in mediating contextual modulation of texture perception, that a discontinuity between target and surround induces segmentation and thus reduces contextual modulation, and that this effect is not observed in our implementation of the feedforward SS model with fixed pooling windows.

### Effect of surround statistics

Besides the geometric cue (the gap) we considered above, another important factor that can reduce contextual modulation is target-surround dissimilarity. The effect of target surround dissimilarity is well reported for object and feature crowding, where the effects of the surround on target identification can be reduced if the two differ in aspects such as color, orientation, or higher-level attributes (Whitney & Levi, 2011; Manassi & Whitney, 2018; Põder, 2007; Louie, Bressler, & Whitney, 2007; Kooi et al., 1994; Farzin, Rivera, & Whitney, 2009), thus increasing target saliency (Gheri, Morgan, & Solomon, 2007). This breakdown in statistical similarity is known to enhance perceptual saliency, (Li, 1999; 2002) and in some cases is suggested to act through segmentation (Whitney & Levi, 2011; Manassi et al., 2013). Understanding the effects of surround structure on contextual modulation of texture perception is important because during natural scene perception there is abundant variability in texture properties and arrangement. Furthermore, different levels of surround structure are often used as proxies for different stages of neural processing (Manassi & Whitney, 2018; Farzin et al., 2009; Louie et al., 2007; Gong, Xuan, Smart, & Olzak, 2018), which could provide insights on the mechanisms behind our observations. For these reasons, we next asked how target-surround dissimilarity affects peripheral texture perception, and how it interacts with segmentation.

#### Experiment 3: FAS dissimilarity but not HOS dissimilarity strongly reduces contextual modulation through segmentation

We focused on target-surround dissimilarity at the FAS and HOS levels because they are related to the SS model (Freeman & Simoncelli, 2011; Freeman et al., 2013) and to physiology (Freeman & Simoncelli, 2011; Freeman et al., 2013; Balas et al., 2009; Ziemba et al., 2016; Okazawa et al., 2015; 2017), as discussed above. Previous work on contextual modulation (Whitney & Levi, 2011; Manassi & Whitney, 2018; Xing & Heeger, 2000) suggests that dissimilar surrounds should have a smaller influence on target perception. Also, it is well known that textures can be segmented from one another based on dissimilarity in their statistics (Victor, Conte, & Chubb, 2017; Rosenholtz, 2014; B. Julesz, 1962), and thus we expect that textures that allow for good target-surround segmentation will lead to reduced contextual modulation. But although effects of FAS and certain HOS in perceptual segmentation and contextual modulation have been studied in a variety of experimental settings (e.g. (Whitney and Levi, 2011; Xing and Heeger, 2000; B. Julesz, 1962; B. Julesz, Gilbert, and Victor, 1978; Victor et al., 2013; Hermundstad et al., 2014; Zavitz and Baker, 2014; Bela Julesz and Caelli, 1979; Victor et al., 2013)), and a wide arrange of computational models attempt to explain texture segmentation and contextual modulation (for reviews and examples see (Victor et al., 2017; Rosenholtz, 2014; Michael S Landy, 2013; Bhatt, Carpenter, and Grossberg, 2007; Axel Thielscher and Neumann, 2005; A. Thielscher, Kölle, Neumann, Spitzer, and Grön, 2008; Bergen and Landy, 1991; Li, 2002)), these processes have not been systematically studied for naturalistic textures, and their effects can also be task dependent (Vancleef et al., 2013; Victor et al., 2017), making it difficult to tell a priori what effects they may have in our task.

To test the effects of FAS and HOS dissimilarity, we compared 3 different surround textures (Figure 6a): 1) The same PS texture as the target (none dissimilar), 2) a different PS texture with FAS and pixel histogram matched to the target PS texture (HOS dissimilar), and 3) a different PS texture with only its pixel histogram matched to the target (FAS & HOS dissimilar). Furthermore, to study the interaction of FAS and HOS dissimilarity with segmentation, we showed these surround textures in both the continuous and discontinuous conditions. In this experiment, target size was the same for the continuous and discontinuous conditions, and the gap was generated by enlarging the surround for the discontinuous condition (increasing inner and outer diameter to maintain its width).

**Figure 6:**
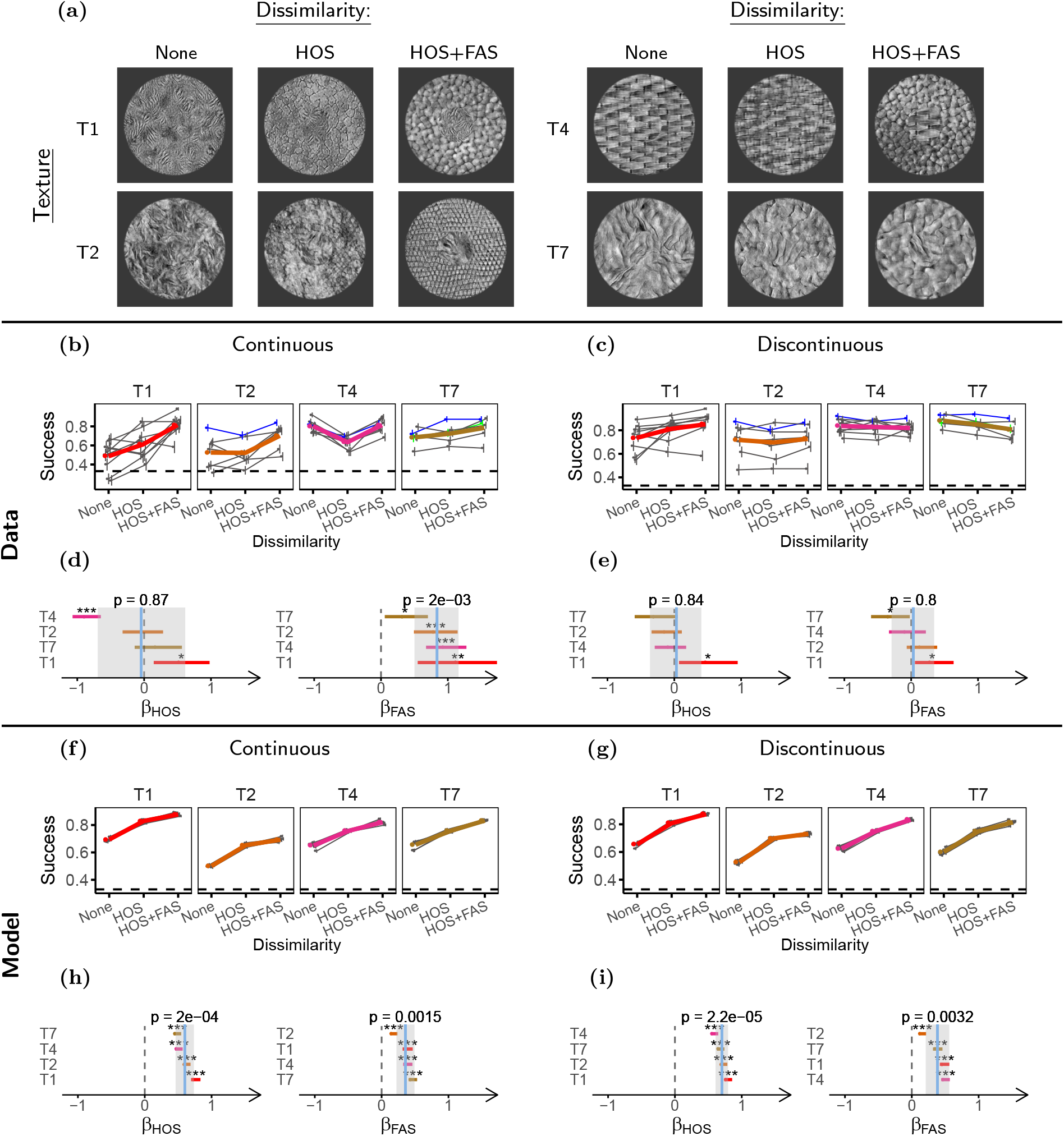
Target-surround dissimilarity reduces contextual modulation. **(a)** Samples of the stimuli used in this experiment, showing for target textures the 3 levels of target-surround dissimilarity used in the experiment (discontinuous stimuli not shown). **(b)** & **(c)** Task performances for the different target-surround dissimilarities in the continuous and discontinuous conditions respectively. **(d)** & **(e)** LOR for HOS (*β_HOS_*) and FAS (*β_FAS_*) dissimilarity in the continuous and discontinuous conditions respectively. **(f)**-**(i)** Same as **(b)**-**(e)** but for the model observers. Participants (n=22) completed 31 experimental sessions, and performed between 60 and 120 trials per condition. Model observers (n=8) discriminated 1500 stimuli per condition. The plots in this figure use the same conventions as the corresponding plots in Figure 5.

We presented participants (n=22) with 4 different target textures (Figure 6a), adding to 31 experimental sessions. To analyze the data we fitted a GLMM with parameters for FAS dissimilarity (*β_FAS_*) and HOS dissimilarity (*β_HOS_*) separately to the continuous and discontinuous conditions, where the effect of FAS dissimilarity is estimated as the change in performance between the condition of HOS dissimilarity and the condition of HOS & FAS dissimilarity.

First, we asked whether the two levels of dissimilarity had an effect for the continuous stimulus. The effect of HOS dissimilarity was close to 0 and not significant (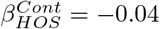, ci = [−0.69,0.61], p = 0.87, Figure 6d), whereas FAS dissimilarity generated strong improvements in performance overall (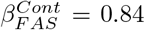, ci = [0.50,1.16], p = 2 × 10^−3^, Figure 6d). We note that the effect of HOS showed considerable variability between textures. In particular, for texture T4 performance was strongly reduced for dissimilar HOS, contrary to expectations. This is likely because the surround without dissimilarity for this texture has a high regularity that introduces a phase effect at the target-surround boundary which could act as a segmentation cue.

To better understand the relation between dissimilarity and segmentation, we then asked whether dissimilarity interacted with discontinuity. If the effects of dissimilarity are mediated simply by surround statistics pooled over fixed regions, we would expect dissimilarity effects for the discontinuous condition comparable to those of the continuous condition (assuming, as we do, a pooling area with the radius of Bouma’s law such that the small change in surround geometry is negligible). On the other hand, if dissimilarity effects are mediated by segmentation, we expect the effects to be reduced in the discontinuous condition where segmentation is already induced by the gap. Consistent with the second mechanism, we found that target-surround dissimilarity had little effect on contextual modulation in the discontinuous condition (Figure 6c) for both HOS (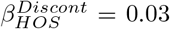, ci = [−0.36,0.40], p = 0.84, Figure 6e), and FAS (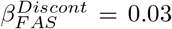, ci = [−0.29,0.34], p = 0.80, Figure 6e), although there was considerable variability between textures. We verified that the change of the effect of FAS dissimilarity for the discontinuous condition was significant (see Supplementary S4 and Figure S6).

Our analysis therefore suggests that FAS dissimilarity effects are strong and mediated by segmentation, whereas HOS dissimilarity effects show considerable variability across textures but are on average weak. We then tested whether these results could be captured by our implementation of the SS model. First, in the continuous condition our model showed a strong improvement in performance when there was HOS dissimilarity, and much weaker changes for FAS dissimilarity (Figures 6f, 6h). Second, these effects were mostly unchanged for the discontinuous condition (Figures 6f, 6h, S6), due to the lack of explicit segmentation processes. Therefore, our implementation of the SS model was not able to capture the patterns observed in the human data.

#### Experiment 4: Naturalistic structure in the surround is important to recruit contextual modulation

The results of the previous section show that a surround with different naturalistic HOS than the target can still exert substantial contextual modulation. Interestingly, other studies have previously shown that contextual modulation can be reduced by removing the natural HOS from the surround. Perceptually, this has been observed for tasks involving recognition and discrimination of natural scenes in peripheral vision (Wallis, Bethge, & Wichmann, 2016; Gong et al., 2018), and for local orientation processing during scene perception (Neri, 2017). Neurally, it has been shown that phase-scrambling the surround (i.e. the HOS are removed but the FAS maintained) strongly affects contextual modulation of neural activity in response to natural images in V1 (Coen-Cagli, Kohn, & Schwartz, 2015; Guo, Robertson, Mahmoodi, & Young, 2005; Pecka, Han, Sader, & Mrsic-Flogel, 2014) and to naturalistic textures in V2 (Ziemba, Freeman, Simoncelli, & Movshon, 2018). This effect of naturalness is thought to reflect that contextual modulation is tuned to natural image statistics, to support efficient coding and optimal perceptual inferences (Coen-Cagli et al., 2015; Pecka et al., 2014). This interpretation seems also in line with previous work with artificial textures, proposing that the asymmetries between textures with uniform and random orientation in texture filling-in could be related to a process of perceptual inference (Hindi Attar, Hamburger, Rosenholtz, Götzl, & Spillmann, 2007). In other work using natural and phase-scrambled scenes, the effect of phase-scrambling has been explained (Gong et al., 2018) as resulting from a weaker engagement of higher areas in the visual hierarchy, leading to reduced contextual modulation in these higher areas. In the context of this literature, our finding of a relatively weak effect of HOS dissimilarity in the previous experiment raises the question of whether the presence of natural HOS is necessary for recruiting contextual modulation for textures.

To address this question, we compared the effects of naturalistic and phase-scrambled surrounds continuous to the target (Figure 7a). Because our experiments required to identify the phase-scrambled target, we reasoned that target-surround similarity with the texture to be identified could affect contextual modulation and lead to unpredictable confounding effects. Therefore, to balance out this possible effect of similarity, we asked half the participants to identify the phase-scrambled texture and the other half to identify the naturalistic texture (modifying the task accordingly, see Methods), and we report the results from both task variants together.

**Figure 7:**
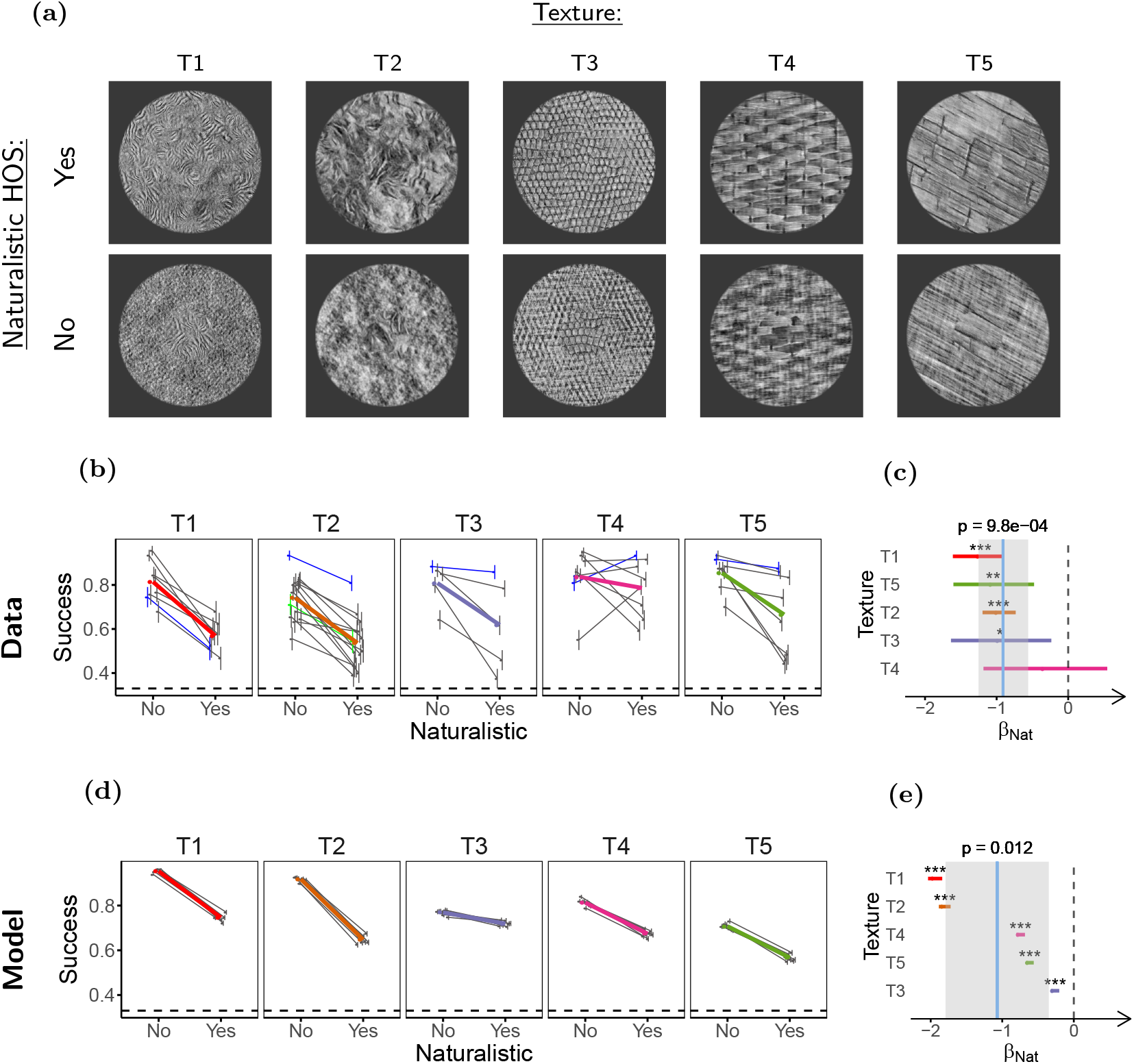
Naturalistic HOS increase contextual modulation. **(a)** Stimuli used in the experiment. Top row: naturalistic surrounds. Bottom row: phase-scrambled surrounds (only naturalistic targets are shown). **(b)** Task performance. **(c)** LOR for the presence of naturalistic HOS (*β_Nat_*). **(d)**, **(e)** Same as **(b)** and **(c)** but for the model observers. Participants (n=43) completed 43 experimental sessions and performed between 90 and 120 trials per condition. Model observers (n=8) discriminated 1500 trials per condition. The plots in this figure use the same conventions as the corresponding plots in Figure 5.

Participants (n=28) were presented with 5 textures, adding to 43 experimental sessions. Consistent with previous studies, we observed that performance was worse with natural HOS in the surround (*β_Nat_* = −0.91, ci = [−1.25, −0.56], p = 10 × 10^−4^, Figure 7c). This is in agreement with previous physiology studies (Coen-Cagli et al., 2015; Guo et al., 2005; Pecka et al., 2014) showing that naturalistic HOS in the surround are important for fully engaging contextual modulation, possibly due to the tuning of contextual modulation to natural image statistics for efficient coding and inference. Together, these results and those from experiment 3 suggests that although the presence of HOS in the surround is important for contextual modulation, their similarity to the HOS of the center is of secondary importance. We note, however, that contextual modulation still occurs for phase-scrambled surrounds (section S5), thus the phenomenon can occurr in the absence of naturalistic HOS.

Although as discussed, the effect of naturalness may reflect the tuning of contextual modulation to natural statistics, (Coen-Cagli et al., 2015; Pecka et al., 2014; Ziemba et al., 2018), we observed a qualitatively similar effect of naturalness in our SS model implementation (Figure 7e). This means that at least part of the effect of naturalness could be mediated by simple pooling. Nonetheless, for textures T1 and T2 we also studied the interaction between naturalness and segmentation (see Supplementary S5, Figure S7), and found that adding a discontinuity reduced the effect of naturalness (*β_Nat:Discont_* = 0.43, ci = [0.06,0.83], p = 0.04, Figure S7d), while this effect was not captured by our SS model (*β_Nat:Discont_* = 0.05, ci = [−0.03,0.12], p = 0.15, Figure S7h). Also, further analysis of the model shows that the observed naturalness effect in the model is not due to surround naturalness itself, but rather due to some specific features of our stimulus generating process (see Supplementary S6, Figure S12). Thus, it is likely that pooling is not the only mediator of the effect of naturalness in our experiments.

In conclusion, these results suggest that naturalistic HOS are important for fully engaging contextual modulation phenomena. This is compatible with suggestions that neuronal contextual modulation phenomena are tuned to the structure of natural images (Coen-Cagli et al., 2015; Pecka et al., 2014), and more specifically, with the results observed for neuronal contextual modulation phenomena in V1 and V2, that may be mechanistically related to our results (see Discussion).

#### Texture crowding

We have thus far shown that texture perception is affected by contextual modulation, and influenced by segmentation and target surround dissimilarity. These characteristics are consistent with a possible role of visual crowding, a contextual modulation phenomenon often regarded as the most important factor of peripheral vision (Rosenholtz, 2016). The SS model explains crowding as a loss of information from pooling together target and surround features when computing local SS (Balas et al., 2009; Freeman & Simoncelli, 2011; Freeman et al., 2013; Whitney & Levi, 2011). However, it is not clear whether this explanation, that is often applied to tasks on non-texture stimuli, should hold for our task. Thus, we decided to test whether the contextual modulation we observed is due to crowding.

There are two main diagnostic criteria for crowding. One is compliance with Bouma’s law, which states that the critical distance at which surrounds interfere with target perception scales linearly with eccentricity with a slope of approximately 0.5 (Pelli et al., 2004). The other is an inwards-outwards asymmetry in which surrounds more eccentric (outwards) to the target exert a stronger modulation than surrounds more central (inwards) to the target (Petrov, Popple, & McKee, 2007; Rosenholtz, 2016; Whitney & Levi, 2011; Farzin et al., 2009; Pelli et al., 2004). Probing Bouma’s law with textures poses experimental challenges, such as changing target-surround distance without breaking continuity or altering target size, and determining how to measure distance between texture stimuli (e.g. (Rosen, Chakravarthi, and Pelli, 2014)). Therefore, we decided to probe the characteristic inwards outwards asymmetry of crowding.

#### Experiment 5: Effect of surround position is small, highly variable and task dependent

To test for inwards-outwards asymmetry in our task, we used half-ring shaped surrounds (Figure 8a) placed inwards or outwards of the target. Participants (n=21) were presented with 5 different textures, completing 37 experimental sessions. Opposite to what has been reported in most crowding studies, performance in our task was consistently lower when the surround was inwards of the target (*β_In_* = −0.32, ci = [−0.47, −0.18], p = 2 × 10^−3^, Figure 8c). This suggests that crowding as reported for classical letter detection or orientation discrimination may not be the main contextual modulation phenomenon in our experiments.

**Figure 8:**
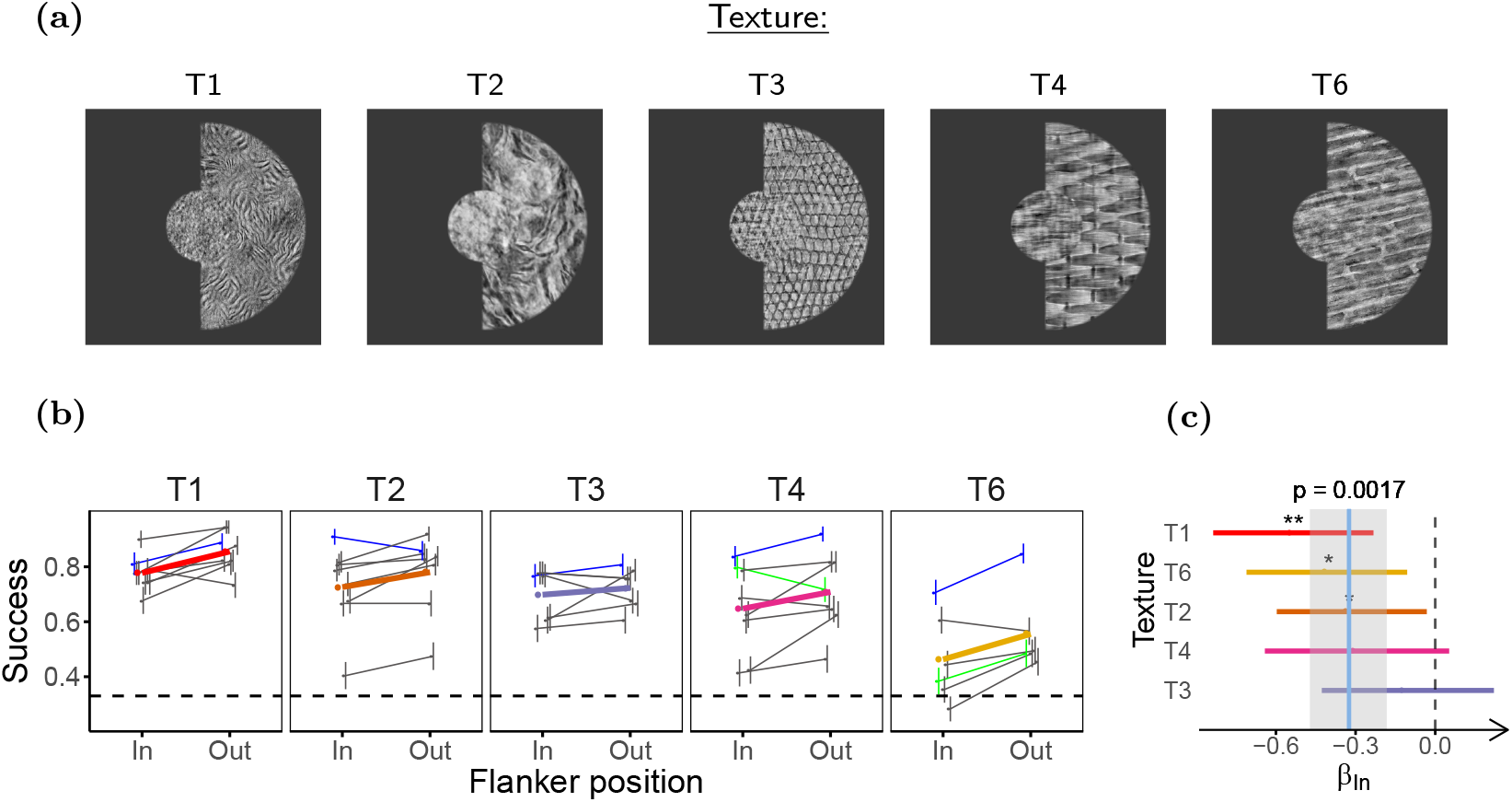
Inwards surrounds affect performance more than outwards surrounds. **(a)** Stimuli used in the experiment. Inwards and outwards surround conditions differ in the position of the half ring of surround texture relative to the fixation point. **(b)** Task performance for the different surround positions. **(c)** LOR for inwards versus outwards surround (*β_In_*). Participants (n=21) completed 40 experimental sessions, and performed between 90 and 99 trials per condition. The figure uses the same conventions as Figure 5.

Nonetheless, unlike the task used here, most reports of inwards-outwards asymmetry use only one target (Petrov et al., 2007; Farzin et al., 2009; Manassi et al., 2012; Banks, Larson, & Prinzmetal, 1979). To verify that the previous result is not only due to this task-related effect, we repeated the experiment for texture T1 using only one target, presented to the right of the fixation point. Participants (n=15) had to report whether the target was naturalistic or phase scrambled. Using this new task we observed an effect of surround position close to 0 (*β_In_* = 0.02, ci = [−0.32,0.38], p = 0.88, Figure 9b). We also verified whether this lack of an effect is due to easier task conditions that bring performance to ceiling levels by using an unsurrounded control condition. Performance was significantly lower for the the surrounded than for the control condition in this experiment (*β_surr_* = −0.46, ci = [−0.72, −0.20], p = 1 × 10^−3^), meaning that the lack of an effect was not due to ceiling performance. This lack of inwards-outwards asymmetry is not what would be expected from the classical asymmetry in crowding, and thus supports the conclusion from the experiment using two targets. Nonetheless, we also note that the difference between the results from the two tasks is in agreement with an effect of task and attention on inwards-outwards asymmetry, such as shown in a previous study in which biasing attention towards the center of the visual field inverted the direction of inwards-outwards asymmetry (Petrov & Meleshkevich, 2011b).

**Figure 9:**
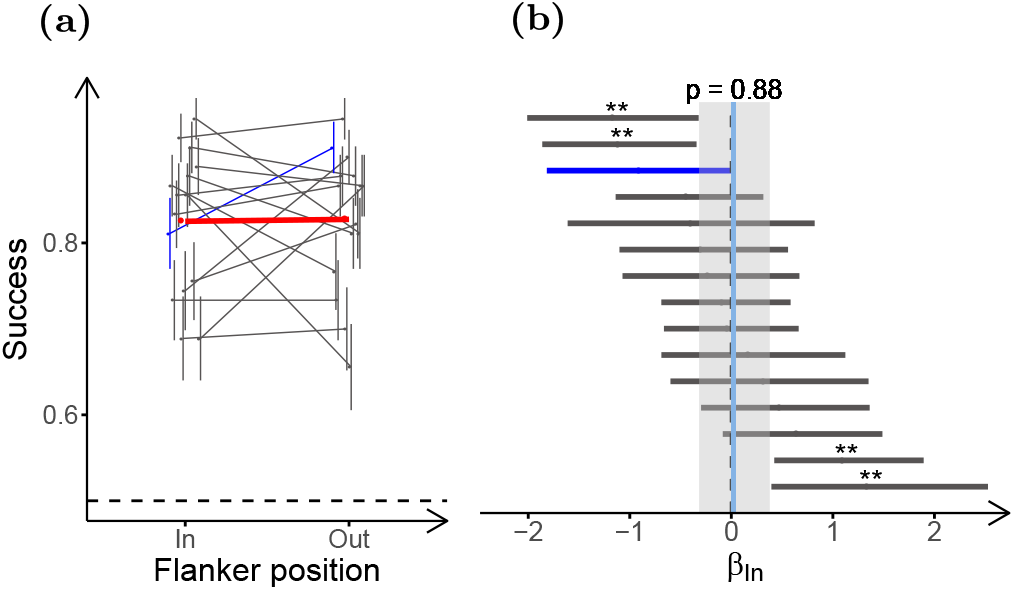
Reduced inwards-outwards asymmetry with single target. **(a)** Task performance for the different surround positions for the task using only one target, for texture T1. **(b)** LOR of the position of the surround for the task using only one target (*β_In_*). Participants (n=15) performed 90 trials per condition. The plots in this figure use the same conventions as the corresponding plots in Figure 3.

Despite the lack of a clear asymmetry in the average performance, variation between participants was high, and some individual participants showed strong effects of surround position in both directions. One plausible interpretation of this result is that contextual modulation in our task arises from different contributing processes, (e.g. crowding and surround suppression, although others processes are possible, see discussion) and that participants with stronger crowding effects would show worse performance for outwards targets, whereas participants more affected by other processes would show little or opposite asymmetry. This hypothesis is in line with previous studies reporting substantial variability in sensitivity to crowding between observers (Lev & Polat, 2015; Petrov & Meleshkevich, 2011a; Wallace, Chiu, Nandy, & Tjan, 2013; Kooi et al., 1994). In addition, we hypothesize that this variability in sensitivity to contextual modulation phenomena could arise from the use of different strategies for solving the task, possibly contributing to the considerable between-participant variability that we observed in the results of the previous experiments.

In conclusion, these results suggest that the processes that underlie crowding in experimental paradigms such as letter recognition, and that have been widely reported to be stronger for outwards surrounds, interact with other processes of at least comparable relevance to contextual modulation of texture perception, that show little or the opposite inwards-outwards asymmetry.

## Discussion

Although the SS model of peripheral vision has had considerable success (Rosenholtz, 2016), studies using complex scenes (Wallis et al., 2019) and simple object-like stimuli (Manassi et al., 2012; 2013; Manassi et al., 2015; Manassi et al., 2016; Doerig et al., 2019; Saarela et al., 2009; Francis et al., 2017) suggest that including processes of segmentation and grouping together with contextual modulation is crucial for a more accurate understanding of peripheral vision. Here we showed that PS texture perception in the periphery is modulated by spatial context, and that contextual modulation is strongly reduced by segmentation engaged both by a gap between target and surround, and by target surround dissimilarity (Figures 4, 5, 6). Although the relevance of segmentation and target-surround dissimilarity for contextual modulation has been studied for discrimination tasks using simple features or objects (Saarela & Herzog, 2009; Manassi et al., 2013; Qiu et al., 2013; Zenger-Landolt & Koch, 2001; Saarela et al., 2010; Kooi et al., 1994; Whitney & Levi, 2011; Manassi & Whitney, 2018; Sayim, Westheimer, & Herzog, 2008), this is to our knowledge the first report of such effects for texture discrimination, which likely involves different processing of the visual input (Jonathan S. Cant, Large, McCall, & Goodale, 2008; J. S. Cant & Xu, 2012; Cavina-Pratesi, Kentridge, Heywood, & Milner, 2010; Rosenholtz, 2014). Furthermore, while the simple feature and object stimuli are more difficult to relate to the SS model (Rosenholtz et al., 2019), our choice of stimuli and task allowed for a direct comparison with the SS model.

In line with previous work using a vernier discrimination task to show that adding more flankers could reduce crowding if these favored target segmentation (Manassi et al., 2012; 2013; Malania, Herzog, & Westheimer, 2007), we found that increasing target size in our texture task can reduce performance if it eliminates a segmentation cue, and that this was not explained by our implementation of the SS model (Figure 4). Also, in line with similar work showing that the precise configuration of the surround is important because it determines grouping with the target (Manassi et al., 2016; Manassi et al., 2013), we show that the precise configuration of the target is important for the same reason. Our SS model implementation was not able to account for this effect (Figure 5). These results thus support the view that the two-stage model with filtering followed by fixed pooling windows cannot fully explain crowding. We note however that this does not argue against the importance of SS as a general framework for understanding peripheral vision, but rather for the need to incorporate segmentation and flexible pooling processes more explicitly. As has been pointed out for previous studies (Rosenholtz et al., 2019), it is possible that a more sophisticated feedforward SS model (e.g. with a nonlinear decoder) could account for some of our segmentation results, leveraging the segmentation cues to extract relevant information from the SS of the stimulus. To test for this, we introduced in experiment 2 and in supplementary section S3 a control for some of the major ways in which this could happen, namely the co-localization of the target and segmentation cue. The small effect of the control gap on task performance of both human participants and our implementation of the SS model suggests that our results cannot be fully explained by an improvement in encoding (or decoding) of target information in the SS of the stimulus facilitated by the low-level properties of the gap. Nonetheless, due to the several changes in geometry introduced in the construction of these control stimuli (Figure 5a), it remains possible that there are some unforeseen changes in the SS of the stimuli that would allow a more elaborate version of the feedforward SS model to account for our results.

Although the effects of different kinds of target-surround similarity on contextual modulation have been widely studied for discrimination tasks using features or objects (see (Manassi & Whitney, 2018; Whitney & Levi, 2011) for reviews), this has not been studied for textures (note that textures have been used to study these effects in the context of contrast perception, e.g. (Solomon, Sperling, and Chubb, 1993; Wang, Heeger, and Landy, 2012)). Our stimulus design allowed us to study the perceptual relevance of dissimilarity in texture properties (specifically, FAS and HOS) to target discrimination. The relation of these properties to the different stages of the SS model and of early visual processing allows us to relate or results to the model and to physiology. Previous studies using artificial textures have reported that FAS is a stronger segmentation cue than HOS, and that some HOS induce moderate and others weak or no segmentation (Zavitz & Baker, 2014; Bela Julesz & Caelli, 1979; Victor et al., 2013). We found that for our naturalistic stimuli dissimilarity in FAS was a strong segmentation cue, but we did not observe clear evidence that dissimilarity in the HOS of the PS model induces segmentation in the periphery (Figure 6). This seems also in agreement with simple inspection of our stimuli, in which the targets strongly pop out when the surround is dissimilar in FAS and HOS, but not when it is only dissimilar in HOS. The weak effect of HOS dissimilarity in peripheral vision is particularly interesting if we note that the textures with HOS dissimilarity were noticeably different under foveal inspection. It is also noteworthy that we did not observe FAS dissimilarity effects when we induced segmentation by a discontinuity between target and surround. If the surround were pooled with the target for the discontinuous condition, as the fixed pooling regions model would suggest, we would expect more similar statistics to interfere more (as was observed for our implementation of the SS model, Figure 6), contrary to what we observed. A possible explanation for this discrepancy between our model and our data is that pooling windows are flexible, and when the surrounds are segmented from the target they are not pooled equally to when grouped together (Wallis et al., 2019; Mareschal, Sceniak, & Shapley, 2001). Finally, we showed that contextual modulation of naturalistic texture perception is strongly dependent on the naturalness of the HOS of the surround (Figure 7), in agreement with previous perceptual (Wallis et al., 2016; Gong et al., 2018; Neri, 2017) and physiological (Coen-Cagli et al., 2015; Guo et al., 2005; Pecka et al., 2014; Ziemba et al., 2018) studies of contextual modulation.

The effects of texture structure may be informative about the mechanism of texture segmentation in the model. Human texture segmentation is a widely studied topic, and several computational models and physiological mechanisms have been proposed in the literature. Our dissimilarity results seem compatible with most of the different existing models, which is not surprising given that they can make similar predictions, and our stimuli were not designed to tell them apart. Nonetheless, our results may offer some interesting constraints on these models, and although an exhaustive analysis is out of the scope of this work, it might be useful to discuss some of the relation to three of the main biologically inspired segmentation models (Michael S Landy, 2013): the feedforward filter-rectify-filter model; the V1-based model with recurrent horizontal connections; and the multi-stage segmentation models with feedback.

In the classic filter-rectify-filter (FRF) kind of segmentation models, texture defined edges are detected by filtering the image with V1-like filters, rectifying the filters outputs, and then applying a second filtering stage on these outputs (Michael S. Landy & Bergen, 1991; Michael S Landy, 2013; Rosenholtz, 2014). The classic version of the model uses a quadratic function for rectification, making it sensitive to local FAS for segmentation, but not to HOS in general (Michael S Landy, 2013), which seems in line with our results. Some models have been proposed to allow the FRF model to be sensitive to some HOS, such as modifying the rectifying function (Zavitz & Baker, 2013) or adding further rectification and filtering steps (Emrith, Chantler, Green, Maloney, & Clarke, 2010), but our results suggest that for naturalistic textures, these further steps may be of secondary importance.

Another class of models compatible with our dissimilarity results are the models based on recurrent contextual interactions at the level of the V1 filtering stage, that lead to differential activation at texture defined edges, allowing for segmentation and saliency (Li, 1999; 2002; Robol, Grassi, & Casco, 2013; Gheorghiu, Kingdom, & Petkov, 2014), which would explain the strong segmentation effect observed for FAS dissimilarity. Interestingly, contextual interactions related to this segmentation model, such as surround suppression and surround normalization, have also been proposed to be a common computation in neural processing (Carandini & Heeger, 2012). If these contextual interactions at the level of V1 are responsible for the FAS-based segmentation, and they are also present in higher areas V2 and V4, we may expect HOS-based segmentation given the selectivity of these areas for the HOS of the PS model (Freeman et al., 2013; Ziemba et al., 2016; Okazawa et al., 2015; 2017). Nonetheless, our results showing weak segmentation for HOS dissimilarities could mean that this process of segmentation may not occur at these higher areas, or that it may be much weaker than in V1 (although see possible limitations below).

The last group of relevant models comprises the more complex and biologically inspired models involving multiple layers and recurrent feedback processing (Bhatt et al., 2007; Axel Thielscher & Neumann, 2005; A. Thielscher et al., 2008; Kim, Linsley*, Thakkar, & Serre, 2019). The complex nature of these models makes them difficult to analyze without actually testing them with our stimuli, although they usually use the first layer of oriented V1-like filters as the substrate of segmentation, allowing for FAS based segmentation. Also, their feedback processing allows them to respond to more complex differences, explaining different texture segmentation results. Our results also provide an interesting experimental test to these models, namely that they should show only weak responses to the HOS explored here.

Besides the results for dissimilarity, it is less clear how our results on naturalness should be related to these models. From the discussion above, it seems that for some of these models, center and surround should not be strongly segmented if they share the same FAS. Nonetheless, it is possible that naturalness effects can emerge in some ways, particularly for the recurrent models. This could be readily tested by using implementations of these models with our stimuli as inputs. Other possible explanations for the effect of naturalness involve segmentation and contextual modulation based on probabilistic inference (Hindi Attar et al., 2007; Coen-Cagli et al., 2015; Pecka et al., 2014), although this would involve at least some extensions on the more mechanistic models described above. Finally, we note that an important limitation of our results is that although the selectivity of areas V2 and V4 to naturalistic HOS is well established (Freeman et al., 2013; Ziemba et al., 2016; Okazawa et al., 2015; 2017), this has not been tested for stimuli with different HOS but matched FAS as those used in this work. Furthermore, the space of PS statistics is high dimensional, and it is possible that other dissimilarities in HOS produce strong segmentation (although note that the textures with dissimilar HOS look considerably different under foveal inspection). Indeed, previous work with artificial textures shows that selectivity for other simpler HOS that can support texture segmentation (Victor et al., 2013) emerges primarily in V2 (Yu, Schmid, & Victor, 2015). Therefore, a more exhaustive exploration of the capacity of naturalistic HOS to induce segmentation would be needed to better understand their role in segmentation, as well as possible contributions from higher visual areas.

What neural mechanisms might underlie the contextual modulation we observe? One candidate is V1 surround suppression, which appears linked to our experimental results in several ways: both strongly depend on FAS similarity (Cavanaugh, Bair, & Movshon, 2002) and on segmentation cues (Coen-Cagli et al., 2015), and it has been proposed that V1 surround suppression underlies perceptual surround suppression (Zenger-Landolt & Heeger, 2003; Carandini & Heeger, 2012), which affects texture perception (Chubb, Sperling, & Solomon, 1989; McDonald & Tadmor, 2006; Wang et al., 2012) and is relatively strong in peripheral vision (Xing & Heeger, 2000; Petrov, Carandini, & McKee, 2005). Also, we showed that contextual modulation of naturalistic texture perception is tuned to the naturalness of the HOS, in agreement with previous perceptual (Wallis et al., 2016; Gong et al., 2018; Neri, 2017) and physiological (Coen-Cagli et al., 2015; Guo et al., 2005; Pecka et al., 2014; Ziemba et al., 2018) studies in contextual modulation. This too could reflect V1 surround suppression, which has been shown to be reduced for scrambled surrounds (i.e. lacking natural HOS) compared to natural images in V1 (Coen-Cagli et al., 2015; Guo et al., 2005; Pecka et al., 2014) (though unpublished recordings indicate this might not be the case for naturalistic textures (Ziemba, Tim Oleskiw, Perez, Simoncelli, & Movshon, 2017)). Overall, our experimental results on contextual modulation and segmentation appear consistent with flexible V1 surround suppression (Coen-Cagli et al., 2015), in which suppression strength is reduced when center and surround are inferred to be segmented on the basis of image statistics. Furthermore, as discussed above, this recurrent process of contextual modulation in V1 is also related to some segmentation models (Li, 1999; 2002; Robol et al., 2013; Gheorghiu et al., 2014; Schmid, 2008), and it could also partly explain the segmentation effects we observed. Following the proposed matching of physiology and the SS model (Figure 1), this process would act after the filtering stage of the model, prior to computing the SS of the texture features.

Another possible mechanism relevant to our results is facilitation. For example, one possible contributor is surround facilitation at the level of V2, observed in texture stimuli similar to ours, in which the response of V2 neurons to a texture patch can be enhanced by naturalistic texture surrounds outside their receptive fields (Ziemba et al., 2018). Following the parallel between physiology and the SS model (Figure 1), this mechanism would act over the output of the SS computation. After the SS of the different image regions are computed, naturalistic surrounds would facilitate the output of the SS computing units corresponding to the target. Although not directly tested in this previous study, this facilitation mechanism could be stronger for scrambled targets than for naturalistic targets, reducing the difference in responses between the two kinds of targets when naturalistic surrounds are included. Thus, this reduced difference between the SS of the two kinds of targets would result in reduced target discriminability. If this is a relevant mechanism in our task, then our results would suggest that V2 surround facilitation is reduced by target surround segmentation, and that it is relatively stronger for phase scrambled targets than for naturalistic targets, which could be readily tested experimentally.

Another mechanism that may contribute to our results is pooling over flexible windows shaped by segmentation, as proposed in studies of natural scene perception in peripheral vision (Wallis et al., 2019) and orientation discrimination in central vision (Mareschal et al., 2001). Flexible surround suppression, facilitation, and flexible pooling windows could therefore be integrated at the corresponding stages of the SS model, leading to a broader framework within which to interpret our results and guide further studies of peripheral vision.

Although our results on contextual modulation and segmentation appeared consistent with perceptual crowding, we did not observe a clear inwards-outwards asymmetry as is often reported for crowding (Petrov et al., 2007) (Figures 8, 9). One possible interpretation for this result is that the processes dominating our contextual modulation effect may not be the same as those in letter crowding. Nonetheless, previous work has shown that the classical inwards-outwards asymmetry can be reversed by biasing attention towards the fovea (Petrov & Meleshkevich, 2011b), and that inward and outwards flankers have different relative weight for different crowding processes and different crowding tasks (Chastain, 1982; Strasburger & Malania, 2013; Strasburger, 2019). Thus, it is possible that our contextual modulation phenomenon is produced by the same mechanisms as letter crowding, but that these mechanisms make different relative contributions to the two phenomena due to the difference in the stimuli used, and that this produces a different overall inwards-outwards asymmetry. Furthermore, we speculate that the large variability we observed across participants in the sensitivity to the different experimental conditions, such as surround position, reflects a variability in the relevance of the different underlying processes (whether the same as in classical crowding or not), which is also consistent with other crowding studies (Lev & Polat, 2015; Petrov & Meleshkevich, 2011a; Wallace et al., 2013; Kooi et al., 1994). To our knowledge, this is the first report of the effects of crowding on texture perception, though crowding has been frequently described as objects undergoing ‘forced texture processing’ (Rosenholtz, 2016). Given that some of the tasks most associated with peripheral vision such as scene perception (Groen, Silson, & Baker, 2017; Ehinger & Rosenholtz, 2016; Brady, Shafer-Skelton, & Alvarez, 2017), guidance of eye movements (Parkhurst & Niebur, 2004; Frey, König, & Einhäuser, 2007; Schmid & Victor, 2014) and the control body of movement (Bardy, Warren, & Kay, 1999; Berencsi, Ishihara, & Imanaka, 2005; Harrington et al., 1985; Sinai, Krebs, Darken, Rowland, & McCarley, 1999; Brandt, Dichgans, & Koenig, 1973) have also been connected to texture perception, further studies on the effects of crowding on texture perception and on these tasks is needed for a complete understanding of peripheral vision limitations in natural vision. Similarly, our results suggest that previous studies of crowding in natural scenes (Wallis & Peter J. Bex, 2012; Gong et al., 2018) might also have measured, to an unknown degree, other contextual modulation processes affecting texture perception. Crowding has been mostly associated with excessive pooling of target and surround features, which can be easily accommodated by the standard SS model. As explained above, our work points to additional processes such as flexible surround suppression and facilitation, highlighting the importance of further studying their relative contributions.

## Acknowledgments

We thank Corey Ziemba for useful discussions on an earlier version of this manuscript. We also thank Adam Kohn for comments on an earlier version of this manuscript. This work was funded by the studentships awarded to D. Herrera and L. Gómez-Sena by the Comisión Académica de Posgrados, UdelaR, Uruguay, and by travelships awarded to D. Herrera by PEDECIBA, UdelaR, Uruguay, and CSIC, Uruguay. Ruben Coen-Cagli was supported in part by National Institutes of Health grant EY031166.

## References

Balas, B., Nakano, L., & Rosenholtz, R. (2009). A summary-statistic representation in peripheral vision explains visual crowding. Journal of Vision, 9(12), 13–13. Publisher: The Association for Research in Vision and Ophthalmology. doi:10.1167/9.12.13

Banks, W. P., Larson, D. W., & Prinzmetal, W. (1979). Asymmetry of visual interference. Perception & Psychophysics, 25(6), 447–456. doi:10.3758/BF03213822

Bardy, B. G., Warren, W. H., & Kay, B. A. (1999). The role of central and peripheral vision in postural control duringwalking. Perception & Psychophysics, 61 (7), 1356–1368. doi:10.3758/BF03206186

Bates, D., Kliegl, R., Vasishth, S., & Baayen, H. (2015). Parsimonious Mixed Models. arXiv:1506.04967 [stat]. arXiv: 1506.04967. Retrieved April 21, 2020, from http://arxiv.org/abs/1506.04967

Bates, D., Martin Maechler, Ben Bolker, Steven Walker, Rune Haubo Bojesen Christensen, Henrik Singmann, … John Fox. (2019). Lme4: Linear Mixed-Effects Models using ‘Eigen’ and S4. Retrieved April 21, 2020, from https://rdrr.io/cran/lme4/

Berencsi, A., Ishihara, M., & Imanaka, K. (2005). The functional role of central and peripheral vision in the control of posture. Human Movement Science. Neural, Cognitive and Dynamic Perspectives of Motor Control, 24 (5), 689–709. doi:10.1016/j.humov.2005.10.014

Bergen, J. R. [James R] & Landy, M. S. [Michael S]. (1991). Computational Modeling of Visual Texture Segregation. In M. Landy & J. A. Movshon (Eds.), Computational Models of Visual Processing (pp. 253–271). MIT Press.

Bhatt, R., Carpenter, G. A., & Grossberg, S. (2007). Texture segregation by visual cortex: Perceptual grouping, attention, and learning. Vision Research, 47(25), 3173–3211. doi:10.1016/j.visres.2007.07.013

Brady, T. F., Shafer-Skelton, A., & Alvarez, G. A. (2017). Global ensemble texture representations are critical to rapid scene perception. Journal of Experimental Psychology: Human Perception and Performance, 43(6), 1160–1176. Place: US Publisher: American Psychological Association. doi:10.1037/xhp0000399

Brandt, T., Dichgans, J., & Koenig, E. (1973). Differential effects of central versus peripheral vision on egocentric and exocentric motion perception. Experimental Brain Research, 16(5), 476–491. doi:10.1007/BF00234474

Burghouts, G. J. & Geusebroek, J.-M. (2009). Material-specific adaptation of color invariant features. Pattern Recognition Letters, 30(3), 306–313. doi:10.1016/j.patrec.2008.10.005

Cant, J. S. [J. S.] & Xu, Y. (2012). Object Ensemble Processing in Human Anterior-Medial Ventral Visual Cortex. Journal of Neuroscience, 32(22), 7685–7700. doi:10.1523/JNEUROSCI.3325-11.2012

Cant, J. S. [Jonathan S.], Large, M.-E., McCall, L., & Goodale, M. A. (2008). Independent Processing of Form, Colour, and Texture in Object Perception: Perception. Publisher: SAGE PublicationsSage UK: London, England. doi:10.1068/p5727

Carandini, M. & Heeger, D. J. [David J.]. (2012). Normalization as a canonical neural computation. Nature Reviews Neuroscience, 13(1), 51–62. Number: 1 Publisher: Nature Publishing Group. doi:10.1038/nrn3136

Cavanaugh, J. R., Bair, W., & Movshon, J. A. (2002). Selectivity and Spatial Distribution of Signals From the Receptive Field Surround in Macaque V1 Neurons. Journal of Neurophysiology, 88(5), 2547–2556. Publisher: American Physiological Society. doi:10.1152/jn.00693.2001

Cavina-Pratesi, C., Kentridge, R. W., Heywood, C. A., & Milner, A. D. (2010). Separate Channels for Processing Form, Texture, and Color: Evidence from fMRI Adaptation and Visual Object Agnosia. Cerebral Cortex, 20(10), 2319–2332. Publisher: Oxford Academic. doi:10.1093/cercor/bhp298

Chastain, G. (1982). Confusability and interference between members of parafoveal letter pairs. Perception & Psychophysics, 32(6), 576–580. doi:10.3758/BF03204213

Chubb, C., Sperling, G., & Solomon, J. A. (1989). Texture interactions determine perceived contrast. Proceedings of the National Academy of Sciences, 86(23), 9631–9635. Publisher: National Academy of Sciences Section: Research Article. doi:10.1073/pnas.86.23.9631

Coen-Cagli, R., Kohn, A., & Schwartz, O. (2015). Flexible gating of contextual influences in natural vision. Nature Neuroscience, 18(11), 1648–1655. Number: 11 Publisher: Nature Publishing Group. doi:10.1038/nn.4128

Cohen, M. A., Dennett, D. C., & Kanwisher, N. (2016). What is the Bandwidth of Perceptual Experience? Trends in Cognitive Sciences, 20(5), 324–335. doi:10.1016/j.tics.2016.03.006

Doerig, A., Bornet, A., Choung, O. H., & Herzog, M. H. (2020). Crowding reveals fundamental differences in local vs. global processing in humans and machines. Vision Research, 167, 39–45. doi:10.1016/j.visres.2019.12.006

Doerig, A., Bornet, A., Rosenholtz, R., Francis, G., Clarke, A. M., & Herzog, M. H. (2019). Beyond Bouma’s window: How to explain global aspects of crowding? PLOS Computational Biology, 15(5), e1006580. Publisher: Public Library of Science. doi:10.1371/journal.pcbi.1006580

Ehinger, K. A. & Rosenholtz, R. (2016). A general account of peripheral encoding also predicts scene perception performance. Journal of Vision, 16(2), 13–13. Publisher: The Association for Research in Vision and Ophthalmology. doi:10.1167/16.2.13

Emrith, K., Chantler, M. J., Green, P. R., Maloney, L. T., & Clarke, A. D. F. (2010). Measuring perceived differences in surface texture due to changes in higher order statistics. JOSA A, 27(5), 1232–1244. Publisher: Optical Society of America. doi:10.1364/JOSAA.27.001232

Farzin, F., Rivera, S. M., & Whitney, D. (2009). Holistic crowding of Mooney faces. Journal of Vision, 9(6), 18–18. Publisher: The Association for Research in Vision and Ophthalmology. doi:10.1167/9.6.18

Francis, G., Manassi, M., & Herzog, M. H. (2017). Neural dynamics of grouping and segmentation explain properties of visual crowding. Psychological Review, 124 (4), 483–504. Place: US Publisher: American Psychological Association. doi:10.1037/rev0000070

Freeman, J. & Simoncelli, E. P. (2011). Metamers of the ventral stream. Nature Neuroscience, 14 (9), 1195–1201. Number: 9 Publisher: Nature Publishing Group. doi:10.1038/nn.2889

Freeman, J., Ziemba, C. M., Heeger, D. J., Simoncelli, E. P., & Movshon, J. A. (2013). A functional and perceptual signature of the second visual area in primates. Nature Neuroscience, 16(7), 974–981. Number: 7 Publisher: Nature Publishing Group. doi:10.1038/nn.3402

Frey, H.-P., König, P., & Einhäuser, W. (2007). The role of first- and second-order stimulus features for human overt attention. Perception & Psychophysics, 69(2), 153–161. doi:10.3758/BF03193738

Friedman, J., Trevor Hastie, Tibshirani, R., Balasubramanian Narasimhan, Noah Simon, & Junyang Qian. (2019). Glmnet: Lasso and Elastic-Net Regularized Generalized Linear Models. Retrieved April 21, 2020, from https://rdrr.io/cran/glmnet/

Gelman, A. & Hill, J. (2006). Data Analysis Using Regression and Multilevel/Hierarchical Models. Google-Books-ID: c9xLKzZWoZ4C. Cambridge University Press.

Gheorghiu, E., Kingdom, F. A. A., & Petkov, N. (2014). Contextual modulation as de-texturizer. Vision Research. The Function of Contextual Modulation, 104, 12–23. doi:10.1016/j.visres.2014.08.013

Gheri, C., Morgan, M. J., & Solomon, J. A. (2007). The Relationship between Search Efficiency and Crowding. Perception, 36(12), 1779–1787. Publisher: SAGE Publications Ltd STM. doi:10.1068/p5595

Gong, M., Xuan, Y., Smart, L. J., & Olzak, L. A. (2018). The extraction of natural scene gist in visual crowding. Scientific Reports, 8(1), 1–13. Number: 1 Publisher: Nature Publishing Group. doi:10.1038/s41598-018-32455-6

Groen, I. I. A., Silson, E. H., & Baker, C. I. (2017). Contributions of low- and high-level properties to neural processing of visual scenes in the human brain. Philosophical Transactions of the Royal Society B: Biological Sciences, 372(1714), 20160102. Publisher: Royal Society. doi:10.1098/rstb.2016.0102

Guo, K., Robertson, R. G., Mahmoodi, S., & Young, M. P. (2005). Centre-surround interactions in response to natural scene stimulation in the primary visual cortex. European Journal of Neuroscience, 21 (2), 536–548. _eprint: https://onlinelibrary.wiley.com/doi/pdf/10.1111/j.1460-9568.2005.03858.x. doi:10.1111/j.1460-9568.2005.03858.x

Harrington, T. L., Harrington, M. K., Quon, D., Atkinson, R., Cairns, R., & Kline, K. (1985). Perception of Orientation of Motion as Affected by Change in Divergence of Texture, Change in Size, and in Velocity. Perceptual and Motor Skills, 61 (3), 875–886. Publisher: SAGE Publications Inc. doi:10.2466/pms.1985.61.3.875

Hermundstad, A. M., Briguglio, J. J., Conte, M. M., Victor, J. D., Balasubramanian, V., & Tkačik, G. (2014). Variance predicts salience in central sensory processing. eLife, 3, e03722. Publisher: eLife Sciences Publications, Ltd. doi:10.7554/eLife.03722

Herzog, M. H., Sayim, B., Chicherov, V., & Manassi, M. (2015). Crowding, grouping, and object recognition: A matter of appearance. Journal of Vision, 15(6), 5–5. Publisher: The Association for Research in Vision and Ophthalmology1. doi:10.1167/15.6.5

Hindi Attar, C., Hamburger, K., Rosenholtz, R., Götzl, H., & Spillmann, L. (2007). Uniform versus random orientation in fading and filling-in. Vision Research, 47(24), 3041–3051. doi:10.1016/j.visres.2007.07.022

John W. Eaton, David Bateman, Soren Hauberg, & Rik Wehbring. (2015). GNU Octave version 4.0.0 manual: A high-level interactive language for numerical computations.

Julesz, B. [B.]. (1962). Visual Pattern Discrimination. IRE Transactions on Information Theory, 8 (2), 84–92. Conference Name: IRE Transactions on Information Theory. doi:10.1109/TIT.1962.1057698

Julesz, B. [B.], Gilbert, E. N., & Victor, J. D. (1978). Visual discrimination of textures with identical third-order statistics. Biological Cybernetics, 31 (3), 137–140. doi:10.1007/BF00336998

Julesz, B. [Bela] & Caelli, T. (1979). On the Limits of Fourier Decompositions in Visual Texture Perception: Perception, 8, 69–73. Publisher: SAGE PublicationsSage UK: London, England. doi:10.1068/p080069

Kim, J., Linsley*, D., Thakkar, K., & Serre, T. (2019). Disentangling neural mechanisms for perceptual grouping. In International Conference on Learning Representations. Retrieved June 26, 2020, from https://openreview.net/forum?id=HJxrVA4FDS

Kleiner, M., David H Brainard, Denis G Pelli, Chris Broussard, Tobias Wolf, & Diederick Niehorster. (2007). What’s new in Psychtoolbox-3? Perception, 36, S14.

Kooi, F. L., Toet, A., Tripathy, S. P., & Levi, D. M. (1994). The effect of similarity and duration on spatial interaction in peripheral vision. Spatial Vision, 8(2), 255–279. Publisher: Brill Section: Spatial Vision. doi:10.1163/156856894X00350

Landy, M. S. [Michael S]. (2013). Texture analysis and perception. In J.S. Werner & L.M. Chalupa (Eds.), The new visual neurosciences (pp. 639–652). MIT Press.

Landy, M. S. [Michael S.] & Bergen, J. R. [James R.]. (1991). Texture segregation and orientation gradient. Vision Research, 31 (4), 679–691. doi:10.1016/0042-6989(91)90009-T

Lazebnik, S., Schmid, C., & Ponce, J. (2005). A sparse texture representation using local affine regions. IEEE Transactions on Pattern Analysis and Machine Intelligence, 27(8), 1265–1278. Conference Name: IEEE Transactions on Pattern Analysis and Machine Intelligence. doi:10.1109/TPAMI.2005.151

Lev, M. & Polat, U. (2015). Space and time in masking and crowding. Journal of Vision, 15(13), 10–10. Publisher: The Association for Research in Vision and Ophthalmology. doi:10.1167/15.13.10

Levi, D. M. (2008). Crowding—An essential bottleneck for object recognition: A mini-review. Vision Research, 48(5), 635–654. doi:10.1016/j.visres.2007.12.009

Li, Z. (1999). Contextual influences in V1 as a basis for pop out and asymmetry in visual search. Proceedings of the National Academy of Sciences, 96(18), 10530–10535. Publisher: National Academy of Sciences Section: Biological Sciences. doi:10.1073/pnas.96.18.10530

Li, Z. (2002). A saliency map in primary visual cortex. Trends in Cognitive Sciences, 6(1), 9–16. Publisher: Elsevier. doi:10.1016/S1364-6613(00)01817-9

Louie, E. G., Bressler, D. W., & Whitney, D. (2007). Holistic crowding: Selective interference between configural representations of faces in crowded scenes. Journal of Vision, 7(2), 24–24. Publisher: The Association for Research in Vision and Ophthalmology. doi:10.1167/7.2.24

Malania, M., Herzog, M. H., & Westheimer, G. (2007). Grouping of contextual elements that affect vernier thresholds. Journal of Vision, 7(2), 1–1. Publisher: The Association for Research in Vision and Ophthalmology. doi:10.1167/7.2.1

Manassi, M., Hermens, F., Francis, G., & Herzog, M. H. (2015). Release of crowding by pattern completion. Journal of Vision, 15(8), 16–16. Publisher: The Association for Research in Vision and Ophthalmology. doi:10.1167/15.8.16

Manassi, M., Lonchampt, S., Clarke, A., & Herzog, M. H. (2016). What crowding can tell us about object representations. Journal of Vision, 16(3), 35–35. Publisher: The Association for Research in Vision and Ophthalmology. doi:10.1167/16.3.35

Manassi, M., Sayim, B., & Herzog, M. H. (2012). Grouping, pooling, and when bigger is better in visual crowding. Journal of Vision, 12(10), 13–13. Publisher:The Association for Research in Vision and Ophthalmology. doi:10.1167/12.10.13

Manassi, M., Sayim, B., & Herzog, M. H. (2013). When crowding of crowding leads to uncrowding. Journal of Vision, 13(13), 10–10. Publisher: The Association for Research in Vision and Ophthalmology. doi:10.1167/13.13.10

Manassi, M. & Whitney, D. (2018). Multi-level Crowding and the Paradox of Object Recognition in Clutter. Current Biology, 28(3), R127–R133. doi:10.1016/j.cub.2017.12.051

Mareschal, I., Sceniak, M. P., & Shapley, R. M. (2001). Contextual influences on orientation discrimination: Binding local and global cues. Vision Research, 41 (15), 1915–1930. doi:10.1016/S0042-6989(01)00082-7

McDermott, J. H. & Simoncelli, E. P. (2011). Sound Texture Perception via Statistics of the Auditory Periphery: Evidence from Sound Synthesis. Neuron, 71 (5), 926–940. doi:10.1016/j.neuron.2011.06.032

McDonald, J. S. & Tadmor, Y. (2006). The perceived contrast of texture patches embedded in natural images. Vision Research, 46(19), 3098–3104. doi:10.1016/j.visres.2006.04.014

McWalter, R. & McDermott, J. H. (2018). Adaptive and Selective Time Averaging of Auditory Scenes. Current Biology, 28(9), 1405–1418.e10. doi:10.1016/j.cub.2018.03.049

Meinecke, C. & Kehrer, L. (1994). Peripheral and foveal segmentation of angle textures. Perception & Psychophysics, 56(3), 326–334. doi:10.3758/BF03209766

Morikawa, K. (2000). Central performance drop in texture segmentation: The role of spatial and temporal factors. Vision Research, 40(25), 3517–3526. doi:10.1016/S0042-6989(00)00170-X

Neri, P. (2017). Object segmentation controls image reconstruction from natural scenes. PLOS Biology, 15(8), e1002611. Publisher: Public Library of Science. doi:10.1371/journal.pbio.1002611

Oberfeld, D. & Stahn, P. (2012). Sequential Grouping Modulates the Effect of Non-Simultaneous Masking on Auditory Intensity Resolution. PLoS ONE, 7(10). doi:10.1371/journal.pone.0048054

Okazawa, G., Tajima, S., & Komatsu, H. (2015). Image statistics underlying natural texture selectivity of neurons in macaque V4. Proceedings of the National Academy of Sciences, 112 (4), E351–E360. doi:10.1073/pnas.1415146112

Okazawa, G., Tajima, S., & Komatsu, H. (2017). Gradual Development of Visual Texture-Selective Properties Between Macaque Areas V2 and V4. Cerebral Cortex, 27(10), 4867–4880. Publisher: Oxford Academic. doi:10.1093/cercor/bhw282

Overvliet, K. E. & Sayim, B. [B.]. (2016). Perceptual grouping determines haptic contextual modulation. Vision Research. Quantitative Approaches in Gestalt Perception, 126, 52–58. doi:10.1016/j.visres.2015.04.016

Paradiso, M. A. & Nakayama, K. (1991). Brightness perception and filling-in. Vision Research, 31 (7), 1221–1236. doi:10.1016/0042-6989(91)90047-9

Parkes, L., Lund, J., Angelucci, A., Solomon, J. A., & Morgan, M. (2001). Compulsory averaging of crowded orientation signals in human vision. Nature Neuroscience, 4 (7), 739–744. Number: 7 Publisher: Nature Publishing Group. doi:10.1038/89532

Parkhurst, D. J. & Niebur, E. (2004). Texture contrast attracts overt visual attention in natural scenes. European Journal of Neuroscience, 19 (3), 783–789. _eprint: https://onlinelibrary.wiley.com/doi/pdf/10.1111/j.0953-816X.2003.03183.x. doi:10.1111/j.0953-816X.2003.03183.x

Pecka, M., Han, Y., Sader, E., & Mrsic-Flogel, T. D. (2014). Experience-Dependent Specialization of Receptive Field Surround for Selective Coding of Natural Scenes. Neuron, 84 (2), 457–469. doi:10.1016/j.neuron.2014.09.010

Pelli, D. G., Palomares, M., & Majaj, N. J. (2004). Crowding is unlike ordinary masking: Distinguishing feature integration from detection. Journal of Vision, 4 (12), 12–12. Publisher: The Association for Research in Vision and Ophthalmology. doi:10.1167/4.12.12

Petrov, Y., Carandini, M., & McKee, S. (2005). Two Distinct Mechanisms of Suppression in Human Vision. Journal of Neuroscience, 25(38), 8704–8707. Publisher: Society for Neuroscience Section: BRIEF COMMUNICATIONS. doi:10.1523/JNEUROSCI.2871-05.2005

Petrov, Y. & Meleshkevich, O. (2011a). Asymmetries and idiosyncratic hot spots in crowding. Vision Research, 51 (10), 1117–1123. doi:10.1016/j.visres.2011.03.001

Petrov, Y. & Meleshkevich, O. (2011b). Locus of spatial attention determines inward–outward anisotropy in crowding. Journal of Vision, 11 (4), 1–1. Publisher: The Association for Research in Vision and Ophthalmology. doi:10.1167/11.4.1

Petrov, Y., Popple, A. V., & McKee, S. P. (2007). Crowding and surround suppression: Not to be confused. Journal of Vision, 7(2), 12–12. Publisher: The Association for Research in Vision and Ophthalmology. doi:10.1167/7.2.12

Põder, E. (2007). Effect of colour pop-out on the recognition of letters in crowding conditions. Psychological Research, 71 (6), 641–645. doi:10.1007/s00426-006-0053-7

Portilla, J. & Simoncelli, E. P. (2000). A Parametric Texture Model Based on Joint Statistics of Complex Wavelet Coefficients. International Journal of Computer Vision, 40(1), 49–70. doi:10.1023/A:1026553619983

Qiu, C., Kersten, D., & Olman, C. A. (2013). Segmentation decreases the magnitude of the tilt illusion. Journal of Vision, 13(13), 19–19. Publisher: The Association for Research in Vision and Ophthalmology. doi:10.1167/13.13.19

R Core Team. (2018). R: A Language and Environment for Statistical Computing. Vienna, Austria.

Riesenhuber, M. & Poggio, T. (1999). Hierarchical models of object recognition in cortex. Nature Neuroscience, 2(11), 1019–1025. Number: 11 Publisher: Nature Publishing Group. doi:10.1038/14819

Robinson, D. & Alex Hayes. (2018). Broom: Convert Statistical Objects into Tidy Tibbles in broom: Convert Statistical Analysis Objects into Tidy Tibbles. Retrieved April 21, 2020, from https://rdrr.io/cran/broom/man/broom.html

Robol, V., Grassi, M., & Casco, C. (2013). Contextual influences in texture-segmentation: Distinct effects from elements along the edge and in the texture-region. Vision Research, 88, 1–8. doi:10.1016/j.visres.2013.05.010

Rosen, S., Chakravarthi, R., & Pelli, D. G. (2014). The Bouma law of crowding, revised: Critical spacing is equal across parts, not objects. Journal of Vision, 14 (6), 10–10. Publisher: The Association for Research in Vision and Ophthalmology. doi:10.1167/14.6.10

Rosenholtz, R. (2014). Texture perception. In The Oxford Handbook of Perceptual Organization. doi:10.1093/oxfordhb/9780199686858.013.058

Rosenholtz, R. (2016). Capabilities and Limitations of Peripheral Vision. Annual Review of Vision Science, 2(1), 437–457. Publisher: Annual Reviews. doi:10.1146/annurev-vision-082114-035733

Rosenholtz, R., Huang, J., Raj, A., Balas, B. J., & Ilie, L. (2012). A summary statistic representation in peripheral vision explains visual search. Journal of Vision, 12(4), 14–14. Publisher: The Association for Research in Vision and Ophthalmology. doi:10.1167/12.4.14

Rosenholtz, R., Yu, D., & Keshvari, S. (2019). Challenges to pooling models of crowding: Implications for visual mechanisms. Journal of Vision, 19(7), 15–15. Publisher: The Association for Research in Vision and Ophthalmology. doi:10.1167/19.7.15

Saarela, T. P. & Herzog, M. H. (2009). Size tuning and contextual modulation of backward contrast masking. Journal of Vision, 9(11), 21–21. Publisher: The Association for Research in Vision and Ophthalmology. doi:10.1167/9.11.21

Saarela, T. P., Sayim, B., Westheimer, G., & Herzog, M. H. (2009). Global stimulus configuration modulates crowding. Journal of Vision, 9 (2), 5–5. Publisher: The Association for Research in Vision and Ophthalmology. doi:10.1167/9.2.5

Saarela, T. P., Westheimer, G., & Herzog, M. H. (2010). The effect of spacing regularity on visual crowding. Journal of Vision, 10(10), 17–17. Publisher: The Association for Research in Vision and Ophthalmology. doi:10.1167/10.10.17

Sayim, B. [Bilge], Westheimer, G., & Herzog, M. H. (2008). Contrast polarity, chromaticity, and stereoscopic depth modulate contextual interactions in vernier acuity. Journal of Vision, 8(8), 12–12. Publisher: The Association for Research in Vision and Ophthalmology. doi:10.1167/8.8.12

Schade, U. & Meinecke, C. (2009). Spatial distance between target and irrelevant patch modulates detection in a texture segmentation task. Spatial Vision, 22(6), 511–527. Publisher: Brill Section: Spatial Vision. doi:10.1163/156856809789822998

Schade, U. & Meinecke, C. (2011). Texture segmentation: Do the processing units on the saliency map increase with eccentricity? Vision Research, 51 (1), 1–12. doi:10.1016/j.visres.2010.09.010

Schmid, A. M. (2008). The processing of feature discontinuities for different cue types in primary visual cortex. Brain Research, 1238, 59–74. doi:10.1016/j.brainres.2008.08.029

Schmid, A. M. & Victor, J. D. (2014). Possible functions of contextual modulations and receptive field nonlinearities: Pop-out and texture segmentation. Vision Research. The Function of Contextual Modulation, 104, 57–67. doi:10.1016/j.visres.2014.07.002

Simoncelli, E., Freeman, W., Adelson, E., & Heeger, D. (1992). Shiftable multiscale transforms. IEEE Transactions on Information Theory, 38 (2), 587–607. Conference Name: IEEE Transactions on Information Theory. doi:10.1109/18.119725

Sinai, M., Krebs, W., Darken, R., Rowland, J., & McCarley, J. (1999). Egocentric Distance Perception in a Virutal Environment Using a Perceptual Matching Task. Proceedings of the Human Factors and Ergonomics Society Annual Meeting, 43(22), 1256–1260. Publisher: SAGE Publications Inc. doi:10.1177/154193129904302219

Solomon, J. A., Sperling, G., & Chubb, C. (1993). The lateral inhibition of perceived contrast is indifferent to on-center/off-center segregation, but specific to orientation. Vision Research, 33(18), 2671–2683. doi:10.1016/0042-6989(93)90227-N

Strasburger, H. (2019). Seven myths on crowding and peripheral vision. PeerJ Preprints. doi:10.7287/peerj.preprints.27353v4

Strasburger, H. & Malania, M. (2013). Source confusion is a major cause of crowding. Journal of Vision, 13(1), 24–24. Publisher: The Association for Research in Vision and Ophthalmology. doi:10.1167/13.1.24

Stürzel, F. & Spillmann, L. (2001). Texture fading correlates with stimulus salience. Vision Research, 41 (23), 2969–2977. doi:10.1016/S0042-6989(01)00172-9

Thielscher, A. [A.], Kölle, M., Neumann, H., Spitzer, M., & Grön, G. (2008). Texture segmentation in human perception: A combined modeling and fMRI study. Neuroscience, 151 (3), 730–736. doi:10.1016/j.neuroscience.2007.11.040

Thielscher, A. [Axel] & Neumann, H. (2005). Neural mechanisms of human texture processing: Texture boundary detection and visual search. Spatial Vision, 18(2), 227–257. doi:10.1163/1568568053320594

Treutwein, B. (1995). Adaptive psychophysical procedures. Vision Research, 35(17), 2503–2522. doi:10.1016/0042-6989(95)00016-X

Vancleef, K., Putzeys, T., Gheorghiu, E., Sassi, M., Machilsen, B., & Wagemans, J. (2013). Spatial arrangement in texture discrimination and texture segregation. i-Perception, 4 (1), 36–52. doi:10.1068/i0515

Venables, W. N. & Ripley, B. D. (2002). Modern Applied Statistics with s (4th ed.). Statistics and Computing. New York: Springer-Verlag. doi:10.1007/978-0-387-21706-2

Vergeer, M. L. T. & van Lier, R. (2007). Grouping Effects in Flash-Induced Perceptual Fading. Perception, 36(7), 1036–1042. Publisher: SAGE Publications Ltd STM. doi:10.1068/p5607

Victor, J. D. (1994). Images, statistics, and textures: Implications of triple correlation uniqueness for texture statistics and the Julesz conjecture: Comment. JOSA A, 11 (5), 1680–1684. Publisher: Optical Society of America. doi:10.1364/JOSAA.11.001680

Victor, J. D., Conte, M. M., & Chubb, C. F. (2017). Textures as Probes of Visual Processing. Annual Review of Vision Science, 3(1), 275–296. doi:10.1146/annurev-vision-102016-061316

Victor, J. D., Thengone, D. J., & Conte, M. M. (2013). Perception of second- and third-order orientation signals and their interactions. Journal of Vision, 13(4), 21–21. Publisher: The Association for Research in Vision and Ophthalmology. doi:10.1167/13.4.21

Wallace, J. M., Chiu, M. K., Nandy, A. S., & Tjan, B. S. (2013). Crowding during restricted and free viewing. Vision Research, 84, 50–59. doi:10.1016/j.visres.2013.03.010

Wallis, T. S. A., Bethge, M., & Wichmann, F. A. (2016). Testing models of peripheral encoding using metamerism in an oddity paradigm. Journal of Vision, 16(2), 4–4. Publisher: The Association for Research in Vision and Ophthalmology. doi:10.1167/16.2.4

Wallis, T. S. A., Funke, C. M., Ecker, A. S., Gatys, L. A., Wichmann, F. A., & Bethge, M. (2019). Image content is more important than Bouma’s Law for scene metamers. eLife, 8, e42512. Publisher: eLife Sciences Publications, Ltd. doi:10.7554/eLife.42512

Wallis, T. S. A. & Peter J. Bex. (2012). Image correlates of crowding in natural scenes. Journal of Vision, 12(6). doi:https://doi.org/10.1167/12.7.6

Wang, H. X., Heeger, D. J., & Landy, M. S. (2012). Responses to second-order texture modulations undergo surround suppression. Vision Research, 62, 192–200. doi:10.1016/j.visres.2012.03.008

Whitney, D. & Levi, D. M. (2011). Visual crowding: A fundamental limit on conscious perception and object recognition. Trends in Cognitive Sciences, 15(4), 160–168. doi:10.1016/j.tics.2011.02.005

Wickham, H. (2016). Ggplot2: Elegant Graphics for Data Analysis. Google-Books-ID: XgFkDAAAQBAJ. Springer.

Wickham, H., Lionel Henry, & RStudio. (2018). Tidyr: Easily Tidy Data with ‘spread()’ and ‘gather()’ Functions.

Wickham, H., Romain François, Lionel Henry, & Kirill Müller. (2018). Dplyr: A grammar of data manipulation.

Willenbockel, V., Sadr, J., Fiset, D., Horne, G. O., Gosselin, F., & Tanaka, J. W. (2010). Controlling low-level image properties: The SHINE toolbox. Behavior Research Methods, 42(3), 671–684. doi:10.3758/BRM.42.3.671

Xie, Y. (2015). Dynamic Documents with r and knitr. Google-Books-ID: lpTYCQAAQBAJ. CRC Press.

Xing, J. & Heeger, D. J. [David J]. (2000). Center-surround interactions in foveal and peripheral vision. Vision Research, 40(22), 3065–3072. doi:10.1016/S0042-6989(00)00152-8

Yu, Y., Schmid, A. M., & Victor, J. D. (2015). Visual processing of informative multipoint correlations arises primarily in V2. eLife, 4, e06604. Publisher: eLife Sciences Publications, Ltd. doi:10.7554/eLife.06604

Zavitz, E. & Baker, C. L. (2013). Texture sparseness, but not local phase structure, impairs second-order segmentation. Vision Research, 91, 45–55. doi:10.1016/j.visres.2013.07.018

Zavitz, E. & Baker, C. L. (2014). Higher order image structure enables boundary segmentation in the absence of luminance or contrast cues. Journal of Vision, 14 (4), 14–14. Publisher: The Association for Research in Vision and Ophthalmology. doi:10.1167/14.4.14

Zenger-Landolt, B. & Heeger, D. J. [David J.]. (2003). Response Suppression in V1 Agrees with Psychophysics of Surround Masking. Journal of Neuroscience, 23(17), 6884–6893. Publisher: Society for Neuroscience Section: Behavioral/Systems/Cognitive. doi:10.1523/JNEUROSCI.23-17-06884.2003

Zenger-Landolt, B. & Koch, C. (2001). Flanker effects in peripheral contrast discrimination—psychophysics and modeling. Vision Research, 41 (27), 3663–3675. doi:10.1016/S0042-6989(01)00175-4

Zhou Wang, A.C. Bovik, H.R. Sheikh, & E.P. Simoncelli. (2004). Image quality assessment: From error visibility to structural similarity. IEEE Transactions on Image Processing, 13(4), 600–612. Conference Name: IEEE Transactions on Image Processing. doi:10.1109/TIP.2003.819861

Ziemba, C. M., Freeman, J., Movshon, J. A., & Simoncelli, E. P. (2016). Selectivity and tolerance for visual texture in macaque V2. Proceedings of the National Academy of Sciences, 113(22), E3140–E3149. Publisher: National Academy of Sciences Section: PNAS Plus. doi:10.1073/pnas.1510847113

Ziemba, C. M., Freeman, J., Simoncelli, E. P., & Movshon, J. A. (2018). Contextual modulation of sensitivity to naturalistic image structure in macaque V2. Journal of Neurophysiology, 120(2), 409–420. Publisher: American Physiological Society. doi:10.1152/jn.00900.2017

Ziemba, C. M., Tim Oleskiw, Perez, R. K., Simoncelli, E. P., & Movshon, J. A. (2017). Selectivity of contextual modulation in macaque V1 and V2. Annual Meeting, Neuroscience. Retrieved April 21, 2020, from https://www.cns.nyu.edu/~lcv/pubs/makeAbs.php?loc=Ziemba17b

