## Supplementary materials for "Flexible contextual modulation of naturalistic texture perception in peripheral vision"

### Supplementary material

#### S1 Converting log-odds ratios to proportion differences

The goal of our statistical analysis is to estimate the effect on performance of changing a given condition, and testing whether the effect is significantly different from 0. All the conditions we analyze can be taken to vary in a binary way, being present or absent (e.g. discontinuity, surround presence, HOS dissimilarity, etc), and thus we frame the analysis as estimating the effect of having that condition present. Also, it is expected that the effect of a given condition can vary across people and across textures. We are therefore interested in estimating the mean effect of that condition across the population of textures and participants, and taking the variability into account when estimating whether it is significantly different from 0.

For this goal we fit a generalized linear mixed model (GLMM) of the binomial family (equivalent to logistic regression) (Gelman & Hill, 2006). This procedure estimates the mean effect of the presence of a given condition A on the probability of success, and also estimates the variability of this effect across the population from which textures and participants were sampled. Taking condition A (e.g. discontinuity) to be a variable with two possible values, A = 1 (A present) and A = 0 (A absent), the model relating A to task performance (probability of success, P) for a given participant $s$ with a given texture $t$ is the following:

$$P(Success|A = a, Texture = t, Participant = s) = \frac{1}{1 + e^{-l(a,t,s)}} \tag{S1}$$

$$l(A = a, Texture = t, Participant = s) = a\beta_A^{t,s} + \beta_0^{t,s} \tag{S2}$$

where $\beta_A^{t,s}$ is the effect of A and $\beta_0^{t,s}$ is the offset for participant s and texture t. Here $\beta_A^{t,s}$ is a sample from a stochastic variable given by $\beta_A^{t,s} = \beta_A + Z_A^t + Z_A^s$. Here $\beta_A$ is the mean effect of condition A (or fixed effect), and $Z_A^t \sim \mathcal{N}(0, \sigma_{A:t}^2)$ and $Z_A^s \sim \mathcal{N}(0, \sigma_{A:s}^2)$ are samples from random variables corresponding to the variability of the effect across textures and participants respectively (the random effects). The random effects are characterized by their standard deviations $\sigma_{A:t}$ and $\sigma_{A:s}$. Similarly, $\beta_0^{t,s} = \beta_0 + Z_0^t + Z_0^s$, with $Z_0^t \sim \mathcal{N}(0, \sigma_{0:t}^2)$ and $Z_0^s \sim \mathcal{N}(0, \sigma_{0:s}^2)$. We note that all participants under a given texture share the same value of $Z_A^t$ and $Z_0^t$, meaning that each texture has its own characteristic effect of A and offset. Fitting a GLMM model to an experiment in which condition A is varied estimates all of the above parameters. The discussions in the text are based on the estimates of the fixed effects for the experimental manipulations, which is the estimated mean effect across textures and subjects.

As mentioned, the effects estimated by the model are the $\beta_A$ coefficients in the linear equation $l$ inside the logistic function. These effects do not directly express the difference in success probabilities between the conditions, they express the log-odds-ratio (LOR) between the conditions, which is a measure related to the difference in probability of success. The odds of success for a given condition can be defined as $odds = \frac{p_{success}}{p_{fail}}$, and it can be converted to probability of success. The odds-ratio is, as the name suggests, the ratio of the odds of two conditions, and it is a measure of the difference in their success probabilities. For example, a way of expressing the effect of condition A on task performance is as the odds-ratio between the condition with A present (A=1) and with A absent (A=0), or $OR_A = \frac{odds_{A=1}}{odds_{A=0}}$. If the presence of A improves performance, $OR_A$ will be larger than 1, and if it hinders performance, $OR_A$ will be between 1 and 0. The log-odds-ratio (LOR) is simply the logarithm of the odds-ratio, and so the parameters estimated by the model can also be understood as $\beta_A = \log OR_A$. The LOR has different advantages as a measure of changes in probability of succes. For one, it is unbounded (can go from $-\infty$ to $+\infty$) and symmetric around 0 (the LOR of a change in probability from $p_1$ to $p_2$ is the opposite of the LOR of a change from $p_2$ to $p_1$). But also, the

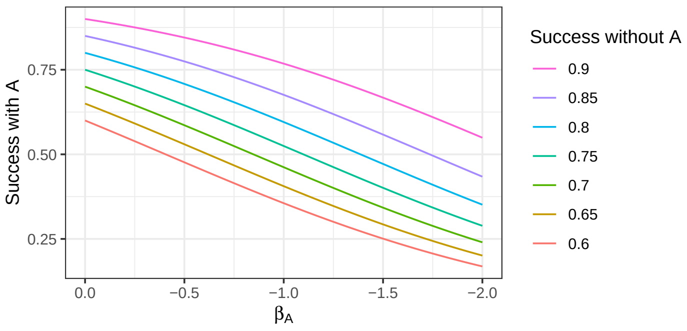


**Figure S1: Reference plot for converting LOR ($\beta_A$) into proportions of correct answers.** Each line indicates a different initial probability of success without condition A (e.g. discontinuity, naturalness, etc). The lines then show the probability of success with condition A for the different values of $\beta_A$.

LOR has the advantage that it reflects an intuitive aspect of the relevance in a change in probability. Intuitively, the relevance of a change in probability depends on the specific probability values, and this is readily reflected in the LOR, but not in the raw probability changes. For example, a change in $p$ from 0.9 to 0.99 is intuitively more important than a change from 0.5 to 0.59. While the change in probability is the same in both cases, the LOR between the conditions are around 2.40 and 0.36 respectively.

But the above mentioned advantage of the LOR means that a given LOR does not uniquely determine a change in probability, since the change in probability will depend on the specific probability values. Therefore we express in the article the estimated effects as the LOR (the $\beta$s). To help get intuition of the magnitude of those effects in terms of probability, Figure S1 shows how to convert LOR to changes in probability. To see how a given LOR translates to a difference in success probability for an experimental manipulation, select an initial probability (a given line) and see what the probability is at the selected LOR. Those would be the probabilities of success without and with condition A present, respectively. Note that a given LOR gives different changes in probability for different initial probabilities.

#### S2 Experiment 1

In experiment 1 we expected the width of the gap to be an important factor in determining the effect of discontinuity because of two opposing factors. For one, since the gap is produced by shrinking the target, a larger gap implies a smaller target for the discontinuous condition. This could lead to a reduction in performance for the discontinuous condition, and potentially mask the effect of segmentation. On the other hand, a smaller gap can be less visible (particularly with the transparency gradients at the borders of the textures), leading to weaker segmentation and thus also leading to a reduced performance for the discontinuous condition. Therefore, we expected the effect of discontinuity to be maximal at some intermediate gap width. We chose to run the experiment using two different gap widths, 0.3° and 0.6°, which seemed to be sufficiently visible but not too large. In experiment 1 we report the result of discontinuity for the gap with 0.3° width. The gap with 0.6° width that showed an effect in the same direction, albeit smaller and non significant ($\beta_{Discont} = 0.27$, ci = $\left[-8 \times 10^{-3}, 0.54\right]$, p = 0.06, Figure S2c). Thus, although the points in the discussion remain unchanged, we note that the experimental effect depends on this relevant parameter.

Also, in experiment 1 we found that texture surrounds impair task performance and that target-

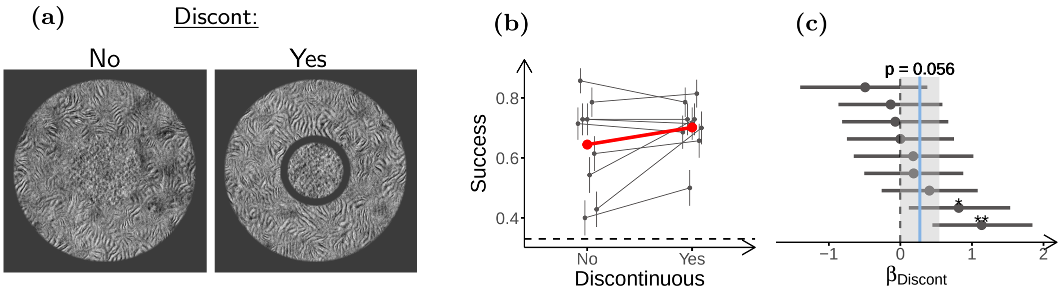


**Figure S2: The effect of discontinuity depends on the size of the gap.** **(a)** Stimulus configurations used in the experiment (only scrambled targets shown). Left: Continuous stimulus (larger target size), Right: Discontinuous stimulus (smaller target size) with a gap of 0.6°. **(b)** Task performance for the two conditions. **(c)** LOR for discontinuity ($\beta_{Discont}$), estimated from the performance data in **(b)**. Participants (n=8) performed 70 trials in each condition. The plots in the figure use the same conventions as Figure 3.

surround segmentation can reduce this impairment. Thus, we tested whether segmentation completely recovered task performance. For this we compared task performance for the unsurrounded target, and for stimuli with surround texture but separated from the targets by a gap (Figure S3a). For the discontinuous condition we pooled the data from the two gap sizes (0.3° and 0.6°). We note that the discontinuous and unsurrounded stimuli had the same target size, only differing in the presence of the discontinuous surround. We found that the discontinuous surrounds still impaired performance ($\beta_{Surr} = -0.40$, ci $= [-0.68, -0.15]$, p $= 5 \times 10^{-3}$), showing that segmentation did not completely remove contextual modulation.

We note that all the conditions mentioned in the section corresponding to experiment 1 (the ones for testing the effect of flanker and the effect of discontinuity), as well as those presented here, were presented in the same experimental session. That is, participants shown in Figures 3 and 4 and in this section are the same individuals (except for a missing participant in Figure 3, which was not presented the unsurrounded due to an error).

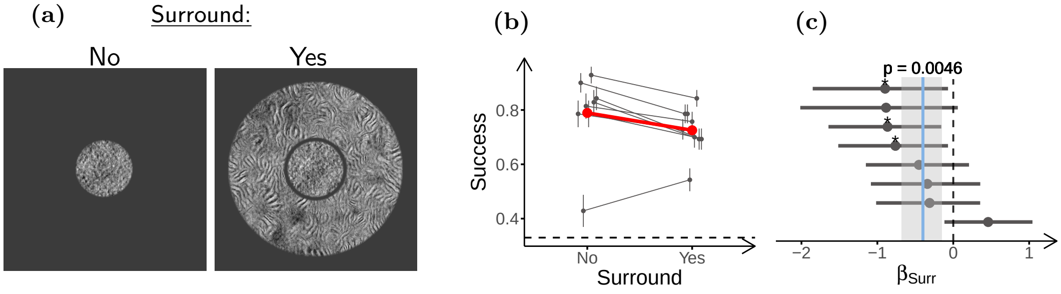


**Figure S3: Discontinuous surrounds generate contextual modulation.** **(a)** Stimulus configurations used in the experiment (only scrambled targets shown). Left: Target without surround, Right: Target with discontinuous surround. **(b)** Task performance for the two conditions. **(c)** LOR for the presence of the discontinuous surround ($\beta_{Surr}$), estimated from the performance data in **(b)**. 8 participants performed 70 trials in the unsurrounded condition, and 140 in the surrounded condition. The plots in the figure use the same conventions as Figure 3.

#### S3 Experiment 2

In experiment 2 we induced continuity by changing target shape from disk-target to split-target (as described in the main text). Although the total area of target texture is roughly the same for the two shapes, it is possible that part of the observed effect could be due to target shape rather than discontinuity. To test for this we included control stimuli for texture T1, consisting of the targets without surrounds, and the targets with surrounds but without a gap, for both target shapes.

First we estimated the effect of target shape by comparing performances for the two targets without surrounds (Figure S4a). The performance for the split-target was slighly worse than for the disk target (Figure S4b), although the effect was small and non-significant ($\beta_{Split} = -0.16$, ci $= [-0.42, 0.11]$, p $= 0.22$, Figure S4c).

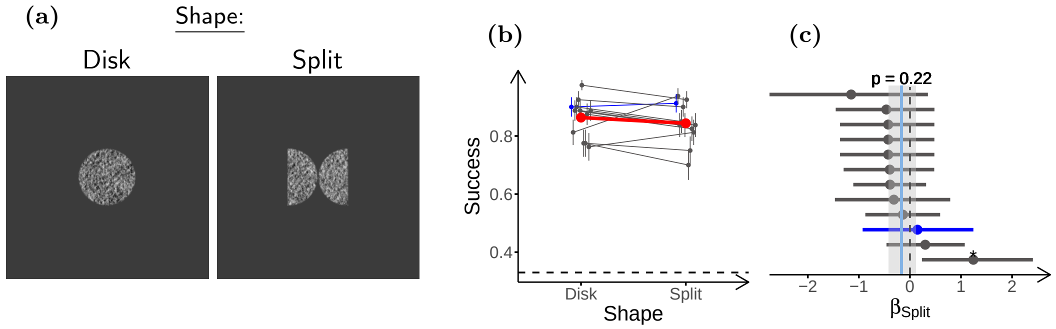


**Figure S4: Target shape does not affect performance for targets without surrounds. (a)** Stimulus configurations used in the experiment. Left: Non-split target (Disk-target) without surround, Right: Split-target without surround. **(b)** Task performance for the two conditions. **(c)** LOR for the splitting of the target ($\beta_{Split}$), estimated from the data in **(b)**. Participants (n=12) performed 80 trials per condition. The plots in the figure use the same conventions as Figure 3.

Then we estimated the effect of target shape and whether it explains the effect of discontinuity. For this we fitted a model with terms for target shape ($\beta_{Split}$), the presence of the gap ($\beta_{Gap}$), and discontinuity ($\beta_{Discont}$) to the stimuli shown in S5a, which include targets of both shapes with surrounds, and in presence and absence of a gap around the target. This analysis also allows to estimate the effect of having a gap around the target after accounting for the discontinuity effect (e.g. effects arising from the low level properties of the gap, or for spatial cueing to the target location within the stimulus). Interestingly, both the effect of target shape ($\beta_{Split} = -0.09$, ci $= [-0.31, 0.14]$, p $= 0.43$, Figure S5c) and of the presence of the gap ($\beta_{Gap} = 0.04$, ci $= [-0.15, 0.22]$, p $= 0.71$, Figure S5d) were close to 0 and non significant, while the effect of discontinuity had a similar magnitude as to that estimated in the main text ($\beta_{Discont} = 0.43$, ci $= [0.14, 0.73]$, p $= 5 \times 10^{-3}$, Figure S5e). This can be seen in the raw performances (Figure S5b), where in the absence of a gap both target shapes have similar performances (thus showing a small effect of target-shape), but in the presence of a gap there is a difference in performance (since one shape is discontinuous with the surround while the other is not). The small effect of the gap can be seen in the little difference between the conditions with and without a gap for the split target, suggesting that the gap does not have a strong effect if it does not induce segmentation. This shows that the improvement in performance described for experiment 2 in the main text comes mainly from discontinuity, rather than target shape or other low level factors that accompany the presence of the gap.

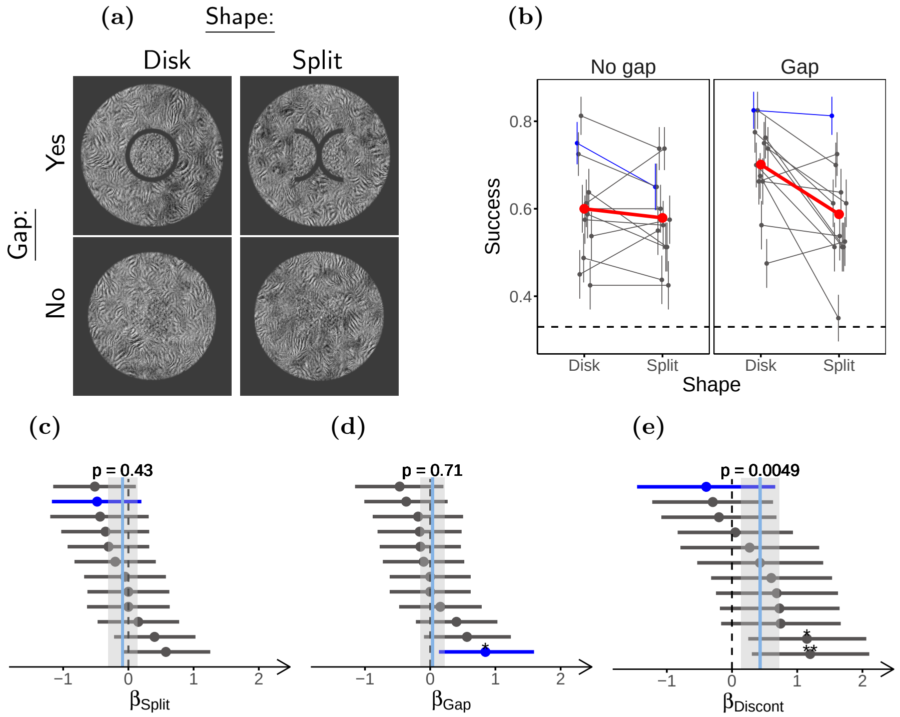


**Figure S5: Target shape does not affect performance for targets with surrounds.** **(a)** Stimulus configurations used in the experiment. Left: Disk-target shape, Right: Split-target shape. Top: Stimuli with a gap. Bottom: Stimuli without a gap. **(b)** Task performance for the four conditions, with the conditions without a gap in the left panel and the conditions with a gap around the target in the right panel. **(c)-(e)** LOR for, respectively, the splitting of the target ($\beta_{Split}$), the presence of the gap ($\beta_{Gap}$), and discontinuity ($\beta_{Discont}$), estimated from the data in **(b)**. Participants (n=12) participants performed 80 trials per condition. The plots in the figure use the same conventions as Figure 3.

#### S4 Experiment 3

In experiment 3, when estimating the effect of target-surround dissimilarity for the discontinuous and continuous conditions separately, there appears to be a considerable change in the effect of FAS dissimilarity. To verify that this change is significant we fitted a GLMM to all the data shown in Figure 6, with parameters for FAS dissimilarity ($\beta_{FAS}$), HOS dissimilarity ($\beta_{HOS}$), discontinuity ($\beta_{Discont}$), and the interactions between discontinuity and dissimilarity ($\beta_{FAS:Discont}$ and $\beta_{HOS:Discont}$). In this model, $\beta_{FAS}$ and $\beta_{HOS}$ are the effect of dissimilarity for the continuous condition, $\beta_{Discont}$ is the effect of discontinuity for the condition with no target-surround dissimilarity, and the interaction terms represent how the effect of dissimilarity changes for the discontinuous condition (or equivalently, how the effect of discontinuity changes for the dissimilar surrounds). We note that due to convergence issues in computing the confidence intervals of the parameters, we excluded the random effects for $\beta_{FAS:Discont}$ from the model, although these were estimated by the model to be close to 0 when fitting the full model, and both the point estimates of the parameters and their corresponding p-values were practically unchanged when including these random effects.

As expected, the full model fit shows a strong and negative effect for the FAS-discontinuity interaction ($\beta_{FAS:Discont} = -0.78$, ci $= [-0.95, -0.61]$, p $= 0.$, Figure S6c) indicating that the effect of FAS dissimilarity is reduced (and practically canceled) when the stimulus is discontinuous. The interaction term between the HOS and discontinuity on the other hand was close to 0 and non significant

($\beta_{HOS:Discont} = 0.07$, ci $= [-0.35, 0.49]$, p $= 0.66$, Figure S6e) and showed considerable variability across textures.

We also estimated the interaction effects for the simulated observers. In line with the similarity between the effects for continuous and discontinuous conditions in Figure 6, the interaction between FAS dissimilarity and discontinuity was close to 0 and non significant ($\beta_{FAS:Discont} = 0.02$, ci $=$ $[-0.02, 0.06]$, p $= 0.38$, Figure S6h). Thus, this confirms the failure of the model to capture the experimental results. Interestingly, the model showed a small but significant interaction between HOS and discontinuity ($\beta_{HOS:Discont} = 0.11$, ci $= [0.07, 0.15]$, p $= 8 \times 10^{-4}$, Figure S6j). This interaction may result from the change in the amount of texture around the target for the discontinuous condition, since the surround texture in the gap is removed, and the magnitude effect will depend on whether HOS are matched or not.

Also, we note that for texture T1 experiment 3 was done using two different widths which, were 0.5° and 1°. Due to the small difference observed between gap sizes in this experiment and preliminary data, we resolved to continue with only the smaller gap for the rest of the experiments. To verify whether the effect of gap width was negligible, we fitted a GLMM to the data with terms for gap width and found them all to be small and non-significant (data not shown). Thus, for the analysis we pooled these two gap sizes together.

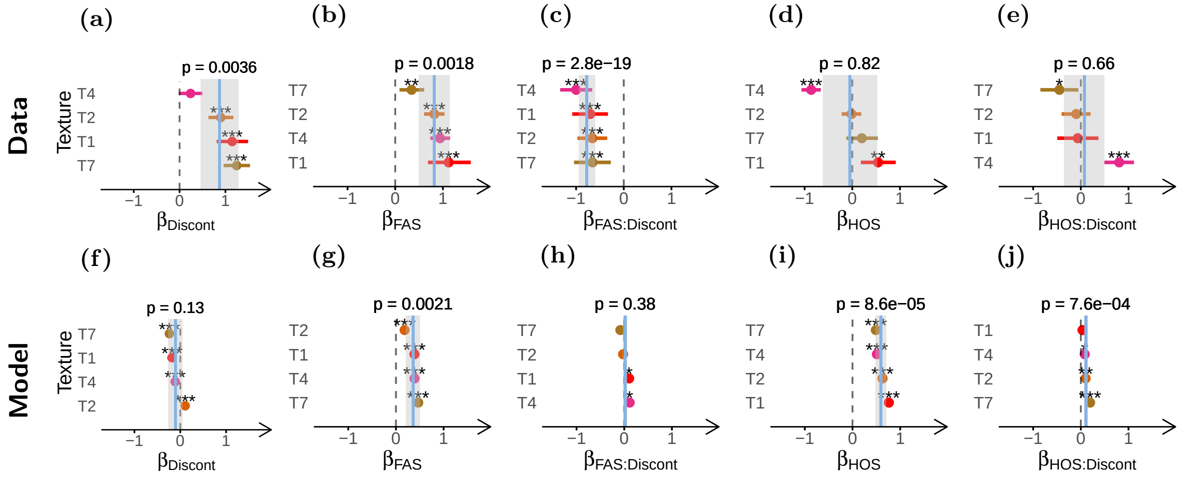


**Figure S6: FAS dissimlarity strongly interacts with discontinuity in participants but not in model observers.** Parameters estimated by fitting a full model to the data from both continuous and discontinuous conditions in experiment 3. **(a)**-**(e)** LOR for, respectively, discontinuity ($\beta_{Discont}$), FAS dissimilarity ($\beta_{FAS}$), discontinuity-FAS interaction ($\beta_{FAS:Discont}$), HOS dissimilarity ($\beta_{HOS}$) and discontinuity-HOS interaction ($\beta_{HOS:Discont}$) estimated from the experimental data. **(f)**-**(j)** Same as **(a)**-**(e)** but for the model observers. The data used for fitting the model is the same as that shown in Figure 6. The plots in the figure use the same conventions as Figure 6.

#### S5 Experiment 4

In the experiment comparing naturalistic and phase scrambled surrounds, we included the discontinuous target-surround conditions for the textures T1 and T2, to determine how segmentation and naturalness interact. We fitted a model to the data with terms for naturalness ($\beta_{Nat}$), discontinuity ($\beta_{Discont}$) and their interaction ($\beta_{Nat:Discont}$). In this model $\beta_{Nat}$ is the effect of naturalness for the continuous condition, $\beta_{Discont}$ is the effect of discontinuity for the phase-scrambled surround, and

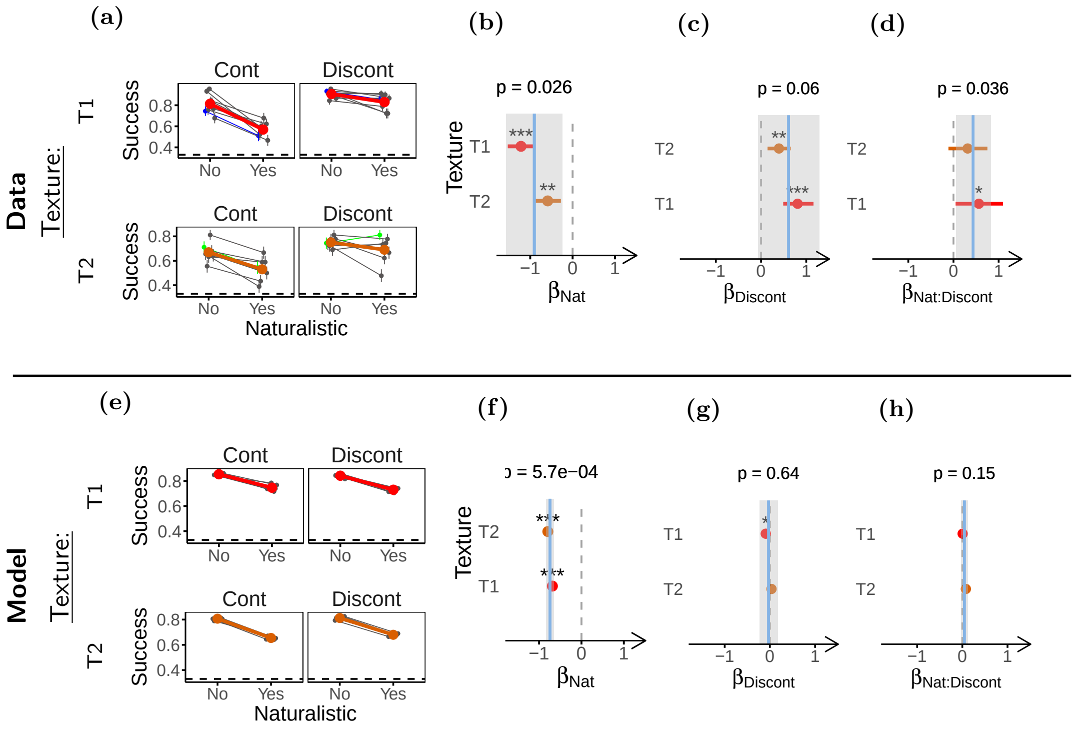


**Figure S7: Surround naturalness interacts with segmentation in participants but not in model observers.** **(a)** Task performance for the naturalistic and phase-scrambled surrounds in both the continuous (left) and discontinuous (right) conditions. Task performances for textures T1 (top) and texture T2 (bottom) are shown. **(b)-(d)** LOR for, respectively, surround naturalness ($\beta_{Nat}$), discontinuity ($\beta_{Discont}$), and their interaction ($\beta_{Nat:Discont}$), estimated from the performance data in **(a)**. **(e)** Same as **(a)** but for the model observers. **(f)-(h)** Same as **(b)-(d)** but for the model observers. Participants (n=28) completed 28 experimental sessions that were included in the analysis, and performed 90 trials per condition.

$\beta_{Nat:Discont}$ represents how the effect of surround naturalness changes for the discontinuous condition (or conversely, how the effect of discontinuity changes when adding surround naturalness). As reported in the main text, there was a strong and significant effect of naturalness for the continuous condition ($\beta_{Nat} = -0.90$, ci $= [-1.57, -0.24]$, p $= 0.03$, Figure S7b). We also observed an effect of discontinuity for the phase-scrambled surround, that did not reach statistical significance for the hierarchical model ($\beta_{Discont} = 0.61$, ci $= [-0.06, 1.30]$, p $= 0.06$, Figure S7c), but did reach significance individually for each texture. Finally, we also observed a moderate and significant interaction between the two terms ($\beta_{Nat:Discont} = 0.43$, ci $= [0.06, 0.83]$, p $= 0.04$, Figure S7d).

While the observer model again captured the effect of naturalness ($\beta_{Nat} = -0.74$, ci $= [-0.83, -0.65]$, p $= 6 \times 10^{-4}$, Figure S7f), it failed to capture its interaction with discontinuity, with an interaction close to 0 and non-significant ($\beta_{Nat:Discont} = 0.05$, ci $= [-0.03, 0.12]$, p $= 0.15$, Figure S7h), thus showing that the effect of naturalness is not fully captured by the feedforward version of the SS model.

Furthermore, for these stimuli we also included the condition with unsurruounded targets in order to determine whether phase-scrambled surrounds produce contextual modulation or not. Performance for the condition with continuous phase-scrambled surrounds was significantly lower than for the unsurrounded targets ($\beta_{Surr} = 0.60$, ci $= [0.18, 1.05]$, p $= 0.02$), showing that naturalistic surrounds are not a necessary condition for contextual modulation.

#### S6 Texture sampling variability in the model

As described in the methods section, we synthesized large textures with the PS algorithm and then, during the task, patches of these textures were randomly sampled on a trial by trial basis to build the stimuli. This procedure introduces some trial by trial variability in the SS of the displayed patches, and thus this constituted a source of stimulus noise in the task. It is not certain what effect this variability could have on the participants performance, since that would likely depend on how they process the stimuli, and on the strategy they use to solve the task. Nonetheless, we propose there are different reasons to think that this noise does not have an important effect on participants. First, participants quickly learned from one or two examples how to perform the task, and they could generalize this to the different texture samples. Second, participant errors could not be predicted from this image variability (analysis not shown). Furthermore, it is very easy to solve the task with foveal inspection, while the task is difficult with limited exposure and peripheral vision. This suggests that the effect of SS variability is probably small as compared to other limitations imposed on the participants by the task design. Lastly, even if the different samples of texture stimuli were always perfectly matched in the PS statistics, it is likely that this will not be the case for the internal representation of SS in participants. This is because different factors such as eye movements, cortical magnification and the use by participants of SS that are not included in the PS model, could still make these different samples unmatched in their internal SS.

On the other hand, given that the implementation of the SS model both lacks the robustness of biological visual systems, and also has few other sources of noise to compete with stimulus variability, the simulated observers may be affected by the sampling variability. Furthermore, understanding the effect of sampling variability on the model can be informative about the behavior of the model, and about possible effects of stimulus noise. Therefore, we performed new simulations removing sampling variability from the task.

In order to remove the sampling variability from the task, we first synthesized for each simulated participant one small PS texture of 128x128 pixels, and its corresponding phase-scrambled texture to be used as targets. Then, to be used for the surrounds, we synthesized one 448x448 texture for each kind of surround texture needed in the experiment to be simulated. All these textures were synthesized as described in the methods, with the exception of their size. Next we used these images to generate one sample of each kind of stimulus (the combination of a kind of surround with a kind of target) using the centermost region of the synthesized textures. This way the crop was not random, and was the same for each stimulus. For example, the naturalistic target was always the exact same patch of texture for the different surrounds, and a given surround was exactly the same for the two kinds of target.

After generating one stimulus of each kind for each simulated subject, we proceeded as in the main simulations to compute the SS and generate pairs of stimuli to discriminate. This results in one kind of stimulus pair for each observer. Finally, we copied the resulting pairs of stimuli to get the needed number of trials to train and test the model (thus there was no variability among samples), and proceeded as described for the main simulations. This way, there was no variability in the stimuli SS in the task (although there was still between trial variability in the noise added to the model).

We note that we used noise with a larger SD than that used in the main simulations. Instead of adding noise with SD equal to that of the SS across the stimuli, we used noise with 10 times that SD. Also, to be able to fit into the synthesized texture for the target, target size was reduced for the first two experiments, using a target size of 100 pixels for experiment 1 and 90 pixels for experiment

2. Also, the plots showing the estimated effects are shown with the same scale as those in the main text, although the effects for this model were considerably smaller. This was done to facilitate the comparison between models and with the experiments.

First, we observed that the performance of the SS model observer was hindered by the presence of the surround in the absence of sampling variability (Figure S8), replicating the results for the original model and the experiments (Figure 3). Then, we repeated experiment 1, by comparing surrounded stimuli with and without a gap. In this case the model had better performance for the discontinuous condition (with gap) than for the continuous (without gap) stimuli (Figure S9), although the effect was very small and non-significant (but it becomes significant when using larger samples). This behavior of the model is the opposite to that observed in the task with stimulus variability (Figure 4), and is opposite from how we expected the SS model to behave, given that the continuous condition has a larger area of informative target texture. One possible explanation for this observation is that, as suggested in previous work (Rosenholtz et al., 2019), strong segmentation cues may allow to more easily recover target SS from the pooled SS, due to factors such as co-localization of a target and a segmentation cue. Despite its small magnitude, this result underscores the unpredictability of the effect of image manipulations on the SS model, which should be particularly problematic for non-texture stimuli as discussed in the main text.

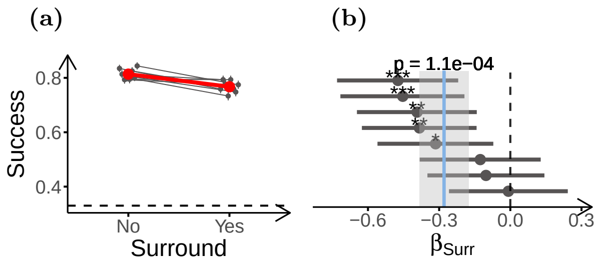


**Figure S8: Performance of the SS model is impaired by surround texture in absence of sampling variability.** **(a)** Task performance of the SS model for stimuli with and without surrounds, in absence of sampling variability. Same layout as Figure 3b. **(b)** LOR for the presence of the surround. Compare to Figure 3c. 8 model observers were tested on 1500 trials per condition.

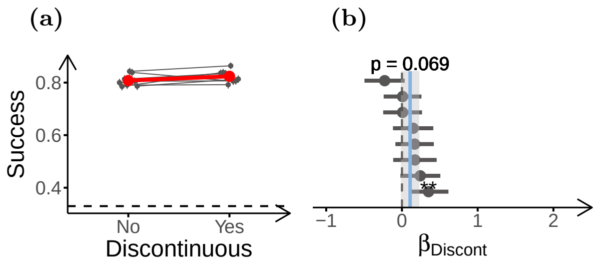


**Figure S9: Performance of the SS model is improved by a discontinuity inducing gap in absence of sampling variability.** **(a)** Task performance of the SS model for the surrounded stimuli with continuous and discontinuous surrounds. Same layout as Figure 3b. **(b)** LOR for discontinuity ($\beta_{Discont}$). Compare to Figure 4c. 8 model observers were tested on 1500 trials per condition.

Although the previous result could raise doubts about the inability of the SS model to explain segmentation by a gap, when testing the effect of discontinuity by changing target shape (experiment 2) we find that model performance is not affected by discontinuity (Figure S10). This result is in agreement with the original simulations, and different from behavior of the participants, which showed better performance for the discontinuous condition (Figure 5). Thus, overall, we still observe that the SS model cannot capture segmentation by a gap.

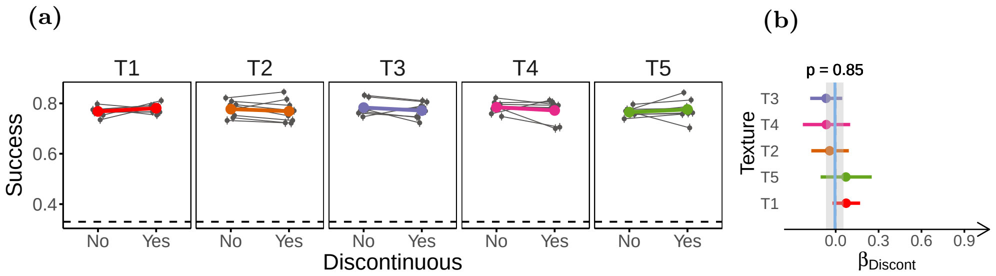


**Figure S10: Performance of the SS model is not affected by discontinuity induced by target-shape in absence of sampling variability.** **(a)** Task performance of the SS model for the surrounded stimuli with continuous and discontinuous surrounds. Same layout as Figure 3b. **(b)** LOR for discontinuity ($\beta_{Discont}$). Compare to Figure 5c. 8 model observers were tested on 1500 trials per condition.

Next we evaluated the behavior of the model for the different target-surround dissimilarities (experiment 3) in absence of variability. The model did not show effects for FAS dissimilarity, or its interaction with discontinuity (Figures S11b, S11c), while it did show a significan albeit small effect for HOS dissimilarity (Figure S11d). This is not an important change as compared to the model with stimulus variability (Figure S6), and therefore does not affect the previous analysis.

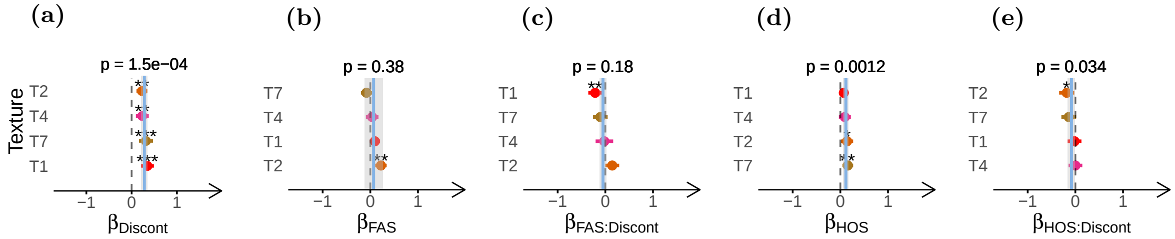


**Figure S11: Performance of the model is improved by target-surround dissimilarity in HOS but not in FAS in absence of sampling variability.** **(a)**, **(b)**, **(c)**, **(d)** & **(e)** LOR for the effects of, respectively, discontinuity ($\beta_{Discont}$), FAS dissimilarity ($\beta_{FAS}$), the interaction between FAS and discontinuity ($\beta_{FAS:Discont}$), HOS dissimilarity ($\beta_{HOS}$), and the interaction between HOS and discontinuity ($\beta_{HOS:Discont}$). Compare to the estimated LOR shown in Figure S6. Same layout as Figure 5c. 8 model observers were tested on 1500 trials per condition.

Lastly, we tested the model on surround naturalness (experiment 4) without stimulus variability. In this experiment sampling variability could be a particularly important factor for the model, because the sampling of the naturalistic images induced a higher variability in the SS than the phase-scrambled images (analysis not shown). In line with this rationale, we observed that the performance of the SS model was not affected by surround naturalness (Figure S12), unlike what was observed in presence of stimulus variability (Figure 7). Thus, whether the SS model can explain the effect of naturalness that was observed for the participants depends on whether the experimental effect is produced by stimulus variability or not. Given that we argued that stimulus variability due to sampling is probably not an important factor for the participants, this means that the SS model may not explain the effect of naturalness in the experimental data.

#### S7 Size adjustment results

A difficulty adjustment procedure was carried out for each individual participant before each experiment (except for texture T1 in experiments 1 and 3), to ensure a similar performance level across

**(a)** **(b)**

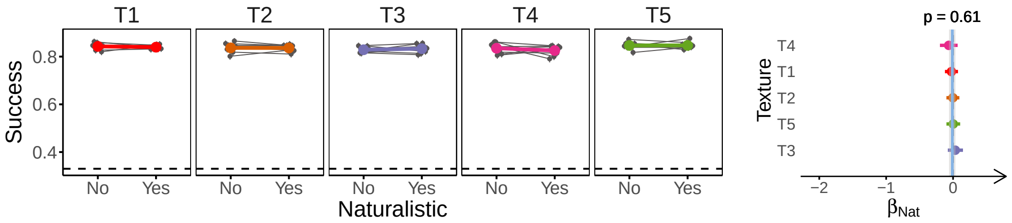


**Figure S12: Performance of the model is not affected by naturalistic HOS in the surround, in absence of sampling variability.** **(a)** Task performance of the SS model for surrounds with the presence or absence of naturalistic HOS. **(b)** LOR for surround naturalness ($\beta_{Nat}$) estimated from the data in **(a)**. Compare to Figure 7. 8 model observers were tested on 1500 trials per condition.

participants. We adjusted target size using the stochastic approximation procedure (Treutwein, 1995). In this procedure, failed responses lead to an increase in target diameter, and successful responses to a decrease, with progressively smaller diameter changes. We set the relation of failure:success step sizes to 9:1, which leads to the procedure converging at a performance of 90% correct responses. The initial target diameter was set between 3.5° and 3.8°, and the initial step size for failed trials was set between 0.7° and 1.0°. The procedure lasted 80 trials or until the step size for the failed trials was smaller than 1 pixel. The actual adaptive procedure started after an initial 10 trials where diameter did not change (totaling 90 trials with the procedure). Figure S13 and Table S1 show the sizes resulting from this adjustment procedure. shows the sizes resulting from this adjustment procedure.

**(a)**

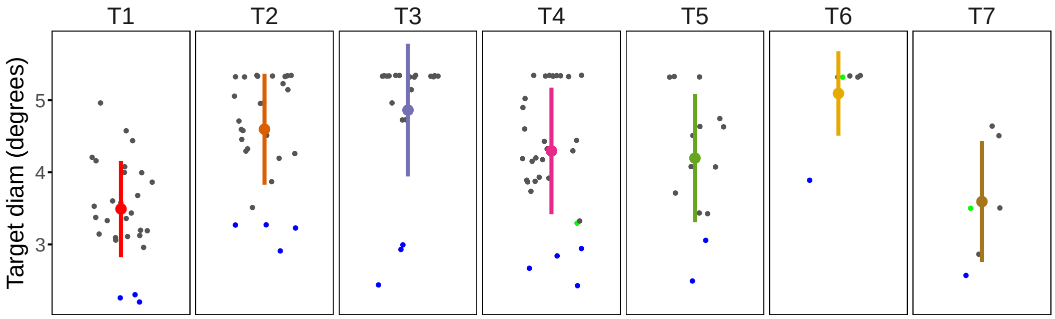


**Figure S13: Target sizes obtained after the difficulty adjustment procedure.** Smaller dots indicate the target size for a given texture session. Blue dots indicate a texture session for author DH and green dots for author LG. Larger colored dots with vertical lines indicate the mean ±SD accross participants. We excluded from the plot the sizes for texture T1 in experiments 1 and 3, in which size was not adjusted, and for the experiment carried out with only one target.

We also tested whether the between participant variability observed in the experimental effects is related to the differences in target size. It is possible that changes in target-surround distances arising from target size adjustment could modify the relative relevance of the contextual modulation processes involved in our task, thus leading to part of the observed between-participant and between-texture variability. We tested this possibility by fitting a linear model to the effect sizes estimated individually for each texture session (that is, for each participant-texture combination, fitting a GLM model only to that texture session data) and the target diameter used in that experimental session. We did this only for experiments 2, 4 and 5 (for the two targets and for the single target experiment, refered to here as 5b), which were the experiments in which only one experimental effect was estimated. For the

| Texture | Mean (deg) | SD (deg) |
|---|---|---|
| T1 | 3.5 | 0.67 |
| T2 | 4.6 | 0.77 |
| T3 | 4.9 | 0.92 |
| T4 | 4.3 | 0.88 |
| T5 | 4.2 | 0.89 |
| T6 | 5.1 | 0.59 |
| T7 | 3.6 | 0.84 |

**Table S1:** Mean and standard deviation of target sizes obtained by the difficulty adjustment procedure for each texture. The sizes from experiments in which the difficulty adjustment was not performed were not taken into account.

experiments with multiple textures (all but 5b) we included a term for texture in the linear model, but removing this term does not change the conclusions from the analysis.

| Experiment | Correlation | p | Df |
|---|---|---|---|
| 2 | 0.25 | $<0.01$ | 34 |
| 4 | 0.02 | 0.61 | 30 |
| 5 | -0.01 | 0.50 | 31 |
| 5b | 0.11 | 0.64 | 13 |

**Table S2:** Pearson correlation coefficients between target diameter and the effect sizes estimated for each texture session in the different experiments.

Table S2 shows the estimated effects of target size on the magnitude of the experimental effect for each experiment. We see that there is a significant effect for experiment 2 but not for the other experiments. The positive sign of the effect in experiment 2 indicates that the larger target size is related to a stronger effect in this experiment. Since most of the experiments did not show an effect, we conclude that target size is not an important factor in explaining between-participant variability.
